# Unravelling Silicon’s Transcriptomic Armor in Soybean against *Macrophomina phaseolina* causing Charcoal Rot Disease

**DOI:** 10.1101/2023.12.22.572986

**Authors:** P. V. Jadhav, S. G. Magar, P. K. Sharma, E. R. Vaidya, M. P. Moharil, S. Jaiswal, S. S. Nichal, R. S. Ghawade, M. S. Iquebal, P. G. Kawar, P. R. Jadhav, S. B. Sakhare, R. B. Ghorade, R. Deshmukh, H. Sonah, D. Kumar, V. K. Kharche, E. A. Torop, R. G. Dani, S. S. Mane

**Author notes:** Correspondence: Pravin V. Jadhav, 1Biotechnology Centre, Department of Agricultural Botany, Dr. Panjabrao Deshmukh Krishi Vidyapeeth, Akola-444104, Maharashtra State, India.

## Abstract

The *Glycine max* L. has been affected by more than 100 diseases, including *Macrophomina phaseolina* producing charcoal rot disease, which reduces production by 70%. In this investigation, RNA-Seq analysis is used for the first time to explore role of silicon in preventing soybean charcoal rot. The study explores the molecular mechanism underlying soybeans’ resilience to charcoal rot when treated with potassium silicon. It was meticulously investigated how *Macrophomina phaseolina* entered the roots. The SEM, which showed a strong link between potassium silicate accumulation and disease resistance. Further investigation indicates that a potassium silicate concentration of 1.7mM lowers disease incidence. Using Illumina HiSeq NGS data, we present a transcriptome analysis revealing genes associated with charcoal rot resistance, highlighting 3,106 genes with distinct expression patterns. The strong enrichment of pathways including “Biosynthesis of ansamycins” and “Flavone and flavonol biosynthesis,” which contribute to resistance against charcoal rot, is highlighted by KEGG enrichment analysis. The ERF transcription factor and NB leucine-rich repeats stands out among the differentially expressed genes as being particularly connected to resistance. The crucial functions that many other important transcription factors, including as MYB, NAC, and proteins from the FAR1 family, play in enhancing soybeans’ resistance to charcoal rot are also noted. This newly discovered information could help in developing tactics to strengthen soybean’s resistance to *Macrophomina phaseolina*.

## 1. Introduction

Soybean (Glycine max L.) holds substantial economic importance due to its protein and oil content. However, its productivity is compromised by various biotic and abiotic stresses [1]. Notably, *Macrophomina phaseolina* causes charcoal rot disease in soybean. In Indian fields, this disease leads to a 77% production reduction [2]. The *Macrophomina phaseolina* is soil-born fungus, produces abundant black microsclerotia, inducing charcoal rot in infected tissue [3]. The fungus moves from roots to stems, obstructing vascular tissue in tap roots [4], and hampers seed germination. In soybean fields, these symptoms emerge post-flowering (R1 stage), peaking during R5, R6, and R7 stages. The *Macrophomina phaseolina* strategies used to the date includes cultural practices, application of fungicide to seeds and biological control, but it has not effective so much or not widely adopted and to control [5]. No plant species has been known to be completely resistant to *Macrophomina phaseolina*, however the soybean genotype DT97-4290 and PI 567562A have been shown to be moderately resistant [6]. Fortunately, Dr. Panjabrao Deshmukh Krishi Vidyapeeth, Akola has created a genotype ‘Suvarn Soya’ (AMS-MB-05-18), resistant to *Macrophomina phaseolina* through mutation breeding in collaboration with the Bhabha Atomic Research Centre, Trombay, India and released at national level during 2019. The host’s molecular responses to the *Macrophomina phaseolina*, however, are not well understood. Determining differentially expressed genes in soybean that are resistant to *Macrophomina phaseolina* is therefore essential. Additionally, silicon (Si) has been employed in several studies to successfully reduce fungi-related diseases in various crops. According to several studies [7], Si deposition in plant cells is associated with processes that increase stress tolerance and stress resistance. Numerous studies demonstrate that exogenous silicon (Si) application enhances resistance against diverse pathogenic fungi impacting both foliar and root health, thereby mitigating fungal infection incidence [8]. Mono-silicic acid, or Si(OH)4, is the type of silicon that plant roots absorb. Si(OH)4 has a poor capacity for soil transport since it is soluble at pH 9 and below 2 mM Si concentration [9]. Therefore, silicon fertilizer can be applied to roots at concentrations up to 2mM of various forms of silicon. The primary sites where silicon is impregnated are the epidermal cell walls of leaf and root cells. According to prior research employing SEM [10], plants may acquire substantial quantities of Si from soil solution but the genetic architecture of silicon response against *Macrophomina phaseolina* has not been deciphered in soybean, yet.

Numerous studies support the use of the transcriptomic (RNA-Seq) approach to pinpoint the molecular evidences of disease resistance [11] & [12]. The rapidly expanding field of transcriptomic has made it feasible to gain a thorough grasp of the intricate connections between plants and their biotic stress. In order to comprehend the fundamental mechanisms behind soybean resistance to *Macrophomina phaseolina* and the impact of silicon treatment, RNA-Seq investigation was proposed. The objective of the present investigation was to ascertain the effects of the charcoal rot disease on soybeans as well as the molecular effectiveness of formulations based on silicon. The knowledge gained from this investigation might be used to develop effective approaches for increasing resistance in soybean in response to the disease known as charcoal rot.

## 2. Results

### 2.1. Molecular characterization of *Macrophomina phaseolina*

*Macrophomina phaseolina* was obtained from diseased soybean plants, and its DNA was extracted from a pure culture utilizing the DNAzol method. The identification of the causative pathogen was accomplished through the employment of the ITS Primer, followed by sequencing of the PCR product. Subsequent analysis of the sequencing data indicated a 98% similarity with *Macrophomina phaseolina*, confirming its molecular identity using the BLAST algorithm. The characterized data was deposited in the NCBI (National Center for Biotechnology Information) database with the Gene Bank accession number MZ823608.1. (https://www.ncbi.nlm.nih.gov/search/all/?term=MZ823608.1)

### 2.2. Field screening

Field screening assessed the phenotypic expression of disease susceptibility and resistance in soybean genotypes under diseased plot conditions. Symptomatic plants were meticulously identified during the V growth stage. The genotype “Suvarn Soya” displayed a disease index ranging from 0 to 1, signifying resistance. While the “TAMS 38” genotype scored 5 on the 0– 5 disease severity scale, indicating its susceptibility (Table 1). Following final confirmation genotypes response to charcoal rot disease, the research transitioned to *in-vitro* screening to evaluate the response of silicon treatment against charcoal rot.

**Table 1:** A disease rating scale for ranking of charcoal rot disease resistant and susceptible genotypes.

| Genotypes | Groups | Disease severity scale | Disease severity index (%) | Disease incidence (%) | Plant mortality |
| --- | --- | --- | --- | --- | --- |
| TAMS 38 | Highly susceptible | 5 (above 75%) | 86 | 94 | 83 |
| Suvarn soya (AMS-MB-05-18) | Resistant | 1 (1-9%) | 4 | 6.5 | 01 |

### 2.3. *In-vitro* screening

*In-vitro* screening was conducted for an RNA sequencing (RNA-seq) study to assess the impact of silicon treatment on the genotype. Soybean plants were inoculated with *Macrophomina phaseolina* in pots, and they were fed with and without 1.7mM Si in three treatment. Both screens on field and *in-vitro* had similar findings, with the Suvarn Soya genotype displaying resistance while the TAMS-38 exhibited susceptibility. The in vitro study involved several small-scale experiments. First, we looked at tissue samples under a microscope (histopathology study). Then, we used SEM and EDX techniques to study how silicon was taken up and accumulated. Finally, we conducted an RNA sequencing (RNA seq) analysis.

#### 2.3.1. Behavior of *Macrophomina phaseolina* under *in-vitro* condition

The histopathological examination provided valuable insights into the crucial stage of the interaction between plants and *Macrophomina phaseolina*. During the study, captivating video footage was obtained, depicting the live movement of the fungus as it initiated the penetration of hyphae into the soybean root and subsequently advanced within the root structure. The video (https://youtube.com/shorts/t1iT5Hic2_M) revealed that fungus *Macrophomina phaseolina* took about 3-4 hours to establish a successful foothold on the soybean root. Histopathological observations of susceptible and resistant genotype roots at different time points post-inoculation (1, 3, 6, 9, and 12 days) unveiled pathogen infiltration within intercellular spaces, causing substantial root tissue damage and plant death. As the infection progressed, notable first observation on the third day taken, only the susceptible genotype exhibited microsclerotia penetration and proliferation, in contrast to the unaffected resistant genotype (Figure 1). Subsequent days saw detrimental effects in the susceptible genotype, including root dehydration and necrotic symptoms by the sixth day in second observation, indicating disease progression. By the ninth day third observation recorded, multiple microsclerotia invasions occurred within the susceptible genotype’s root tissue, leading to browning. In stark contrast, the resistant genotype Suvarn soya had a clear area that was infected with only a few microsclerotia invading. The big difference between the two genotypes became really obvious by the fourth observation on the twelfth day. At that point, the inner tissue of the susceptible genotype was completely surrounded by microsclerotia, covering the parts responsible for transporting water and nutrients. Remarkably, the resistant plant remained unaffected by the infection (Figure 1A-E). These findings underscore the susceptibility of the susceptible genotype to disease, emphasizing the role of hyphae and microsclerotia formation in *Macrophomina phaseolina* pathogenicity.

**Figure 1:**
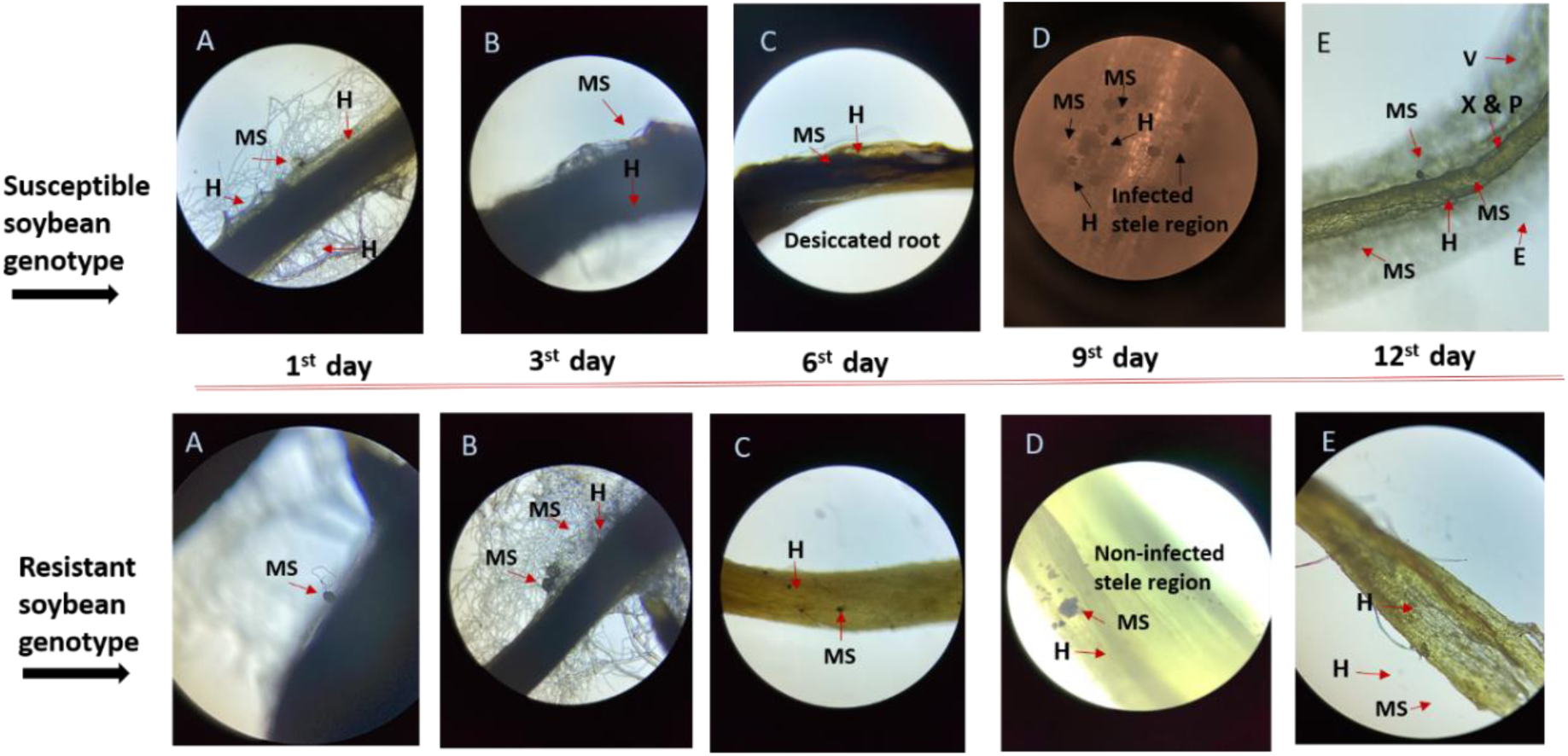
Average trend of 3 days’ interval progress of Charcoal rot disease in soybean genotypes. A) Microsclerotia germination B) Infection and hyphal penetration C-E) Pathogen colonization into infected inner tissues (MS: Microsclerotia; H: Hyphae; X & P: Xylem and Phloem; V: Vessels; E: Epidermis;

### 2.4. Effect of Silicon treatment

Experimental treatments were fed with and without 1.7mM K2SiO3 to compare the effects of silicon against charcoal rot disease. Significant variations between the susceptible and resistant genotypes were recoded for each treatment (Figure 2A). The silicon treated plants grew stronger and revealed that the treated plants were healthier than the untreated ones. We cut the lower stem of plants in half and looked at it under a microscope. However, when we treated the plant with K2SiO3 at a concentration of 1.7mM, the infection was stopped. On the other hand, the stem of a diseased plant that wasn’t treated had a lot of sclerotia and turned black (Figure 2B). Comparing treated and untreated seeds from a susceptible plant, observed a significant difference. Infected plant seeds look bad with blackening, rot, and lower weight. But the silicon-treated plant seeds look healthy and normal, like the uncontaminated ones. (Figure 2C). The key observation noted in the susceptible plants, silicon treatments reduced fungal infection, prevented wilting, and eliminated the fungus, proving their effectiveness against charcoal rot disease. Additionally, silicon accumulation was studied using SEM, showing that as silicon deposition increased, disease incidence decreased.

**Figure 2:**
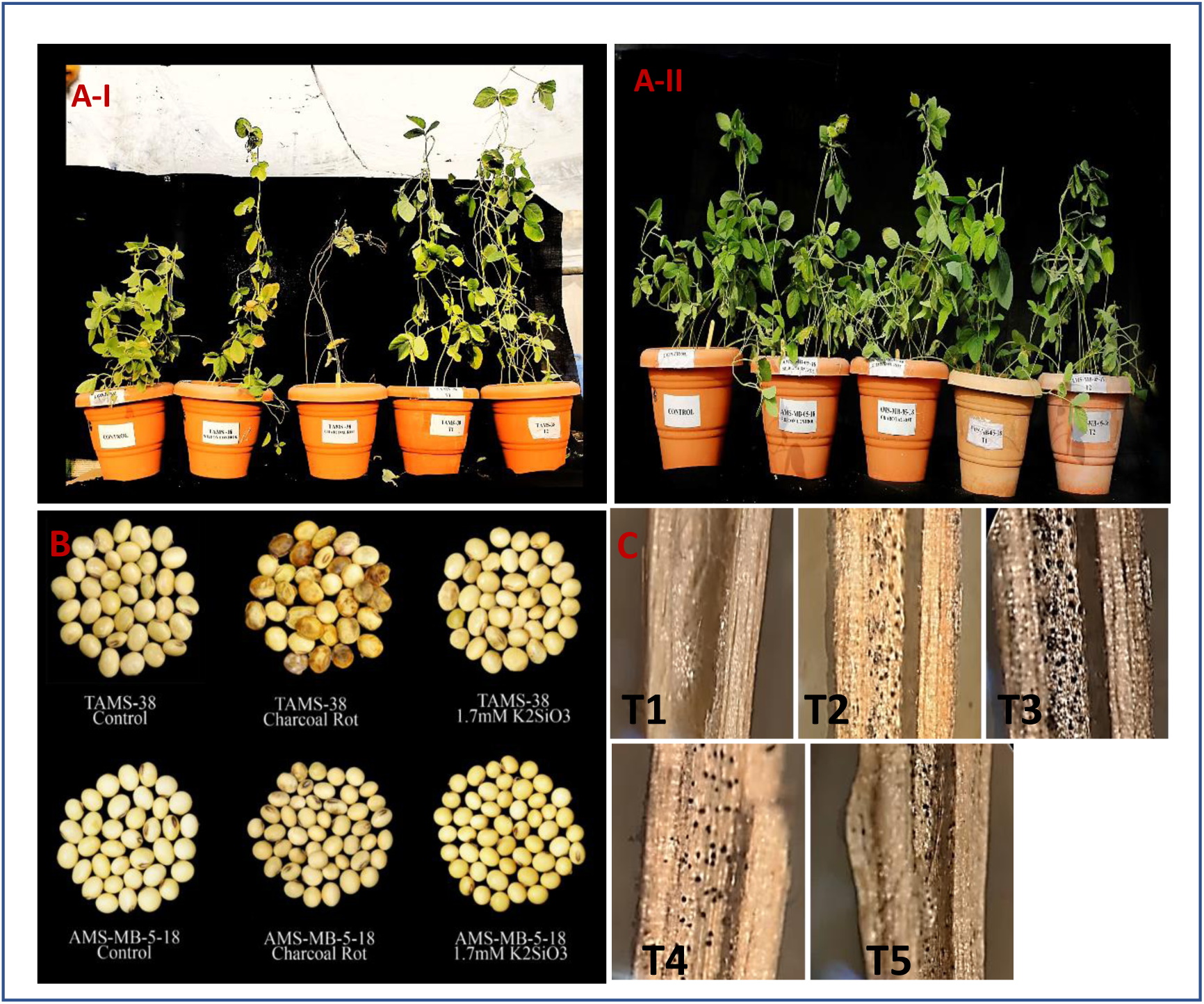
*in-vitro* screening for charcoal rot disease severity. (A) Comparative *in-vitro* screening of soybean genotypes A-I) susceptible genotype (TAMS-38) A-II) resistant genotype (AMS-MB-05-18) for Si effect against charcoal rot. T1: Control; T2: Silicon control; T3: Charcoal rot; T4: 0.7 mM Potassium silicate; T5: 1.7 mM Potassium silicate. (B): Symptoms of charcoal rot showing presence of fungal sclerotia in T1 to T5 of TAMS-38 susceptible genotype charcoal coloured growth observed in infected soybean stem prominently in T3 whereas, T4 & T5 observed with minor infection of C.R. (C): Seeds development in susceptible and resistant soybean genotypes with treatment and without treatment.

### 2.5. Scanning Electron Microscopy of accumulation of silicon in leaves and roots

SEM was used to identify the silicon deposition in the trichomes of leaves and the vascular tissue of roots. Red hue in the trichomes of leaves indicates an accumulation of silicon (Figure 3 A&B). The larger silicon particles found lodged in the roots’ vascular tissue. Resistance was noticed to rise as silicon deposition increased in the root and leaves. Expanding on silicon deposition findings, the study used RNA-seq for transcriptomic analysis to understand how silicon treatment controls the response to *Macrophomina phaseolina* infection in different genotypes. Six samples were collected, consisting of control samples, infected samples, and samples treated with 1.7 mM K2SiO3 for both genotypes. These samples were collected at 42 days after planting, providing a snapshot of gene expression profiles at a critical stage.

**Figure 3:**
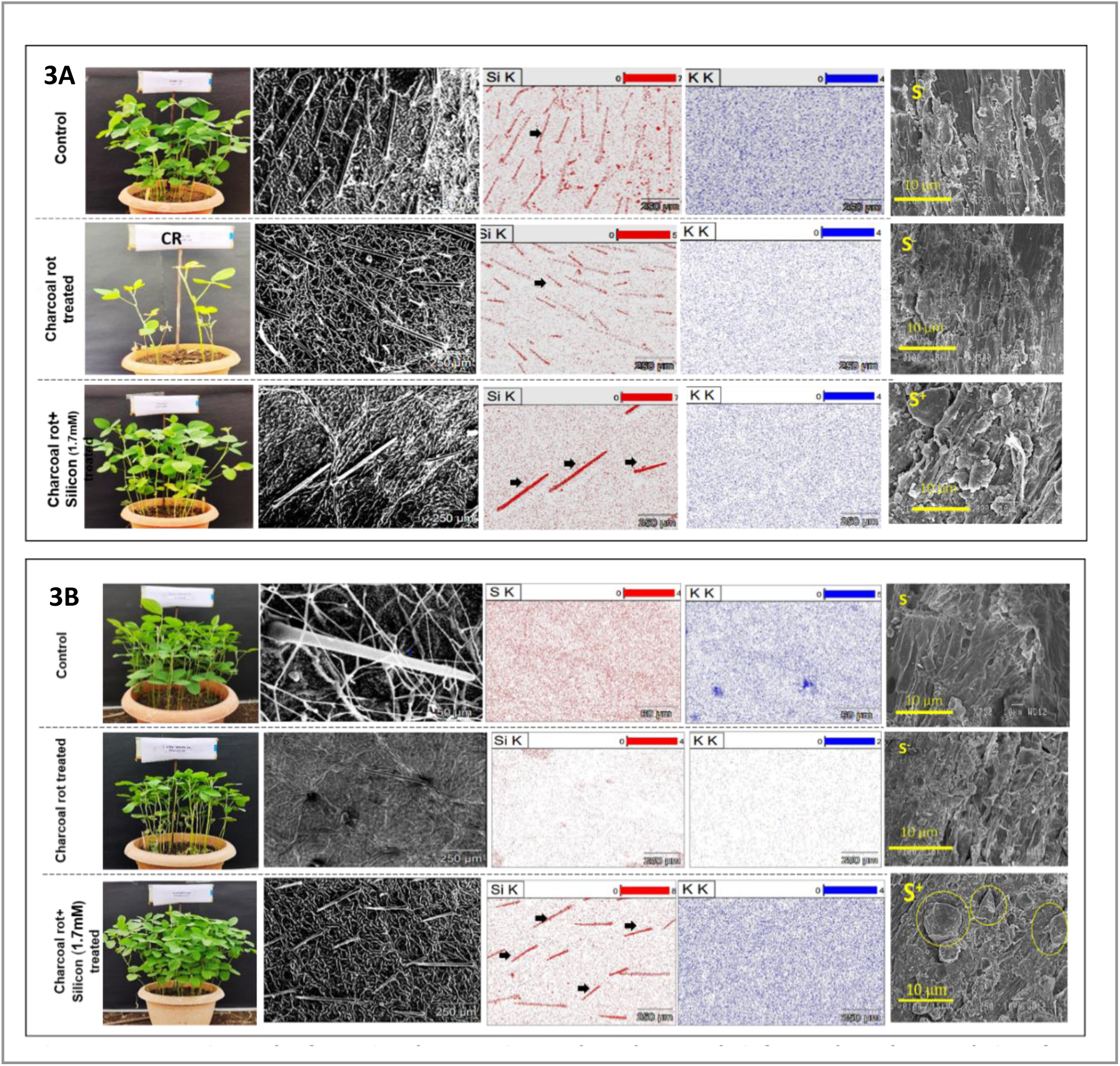
Comparative study of SEM and EDX analysis for uptake and accumulation of silicon in soybean genotype cv. (A)TAMS-38, (B): AMS-MB-5-18, tissue leaf and root.

### 2.6. Raw data Pre-processing and transcriptome assembly

For each of the six-leaf tissue sample from the two dissimilar soybean types, sequencing data were received. A 150-base pair read was used. 16,933,056 read pairs were obtained for the susceptible control, 17,224,957 for the susceptible infected, and 20,157,934 for the susceptible treated. By contrast, 17,666,231 read pairs were obtained for the resistant control, 17,678,885 for the resistant infected, and 23,959,344 read pairs were obtained for the resistant treated. Summary of reads before and after trimming is presented in table 2. The data was of good quality, and 102,074,449 high-quality reads from all the samples were pooled for reference and de-novo-based transcriptome assembly following reads’ pre-processing and contamination removal. A total of 227,790 genes and 410,789 transcripts were produced by the Trinity assembler. The average contig length was determined to be 999.38 base pairs, with a GC content of 40.53%, and the N50 value was 1894 base pairs. There are a total of 117,311 transcripts with a N50 value of 2.699 Kb in the case of reference-based assembly. Table 3 offers a comparison visualization of both assemblies. A BLAST+ comparison of the two assembles showed that they had 110173 transcripts in common, with the de-novo assembly having 48.36% of the total and the reference-based assembly having 94.12%, proving the inclusion of unique transcripts in the de-novo assembly. There were 117617 new transcripts in the de-novo-based assembly and 6880 in the reference-based assembly (Figure 4).

**Figure 4.**
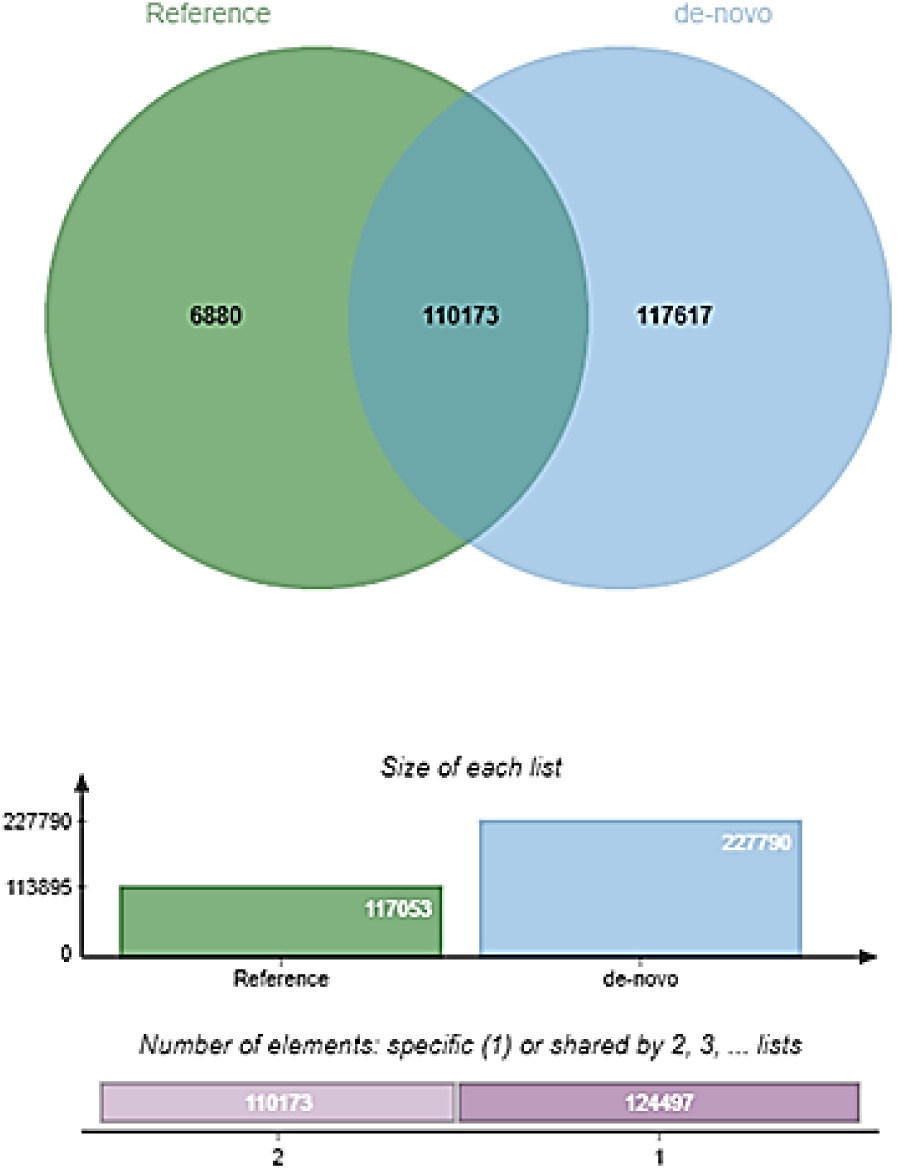
Comparison of reference and *de-novo-*based assemblies in *Macrophomina phaseolina*.

**Table 2:** Summary of Reads’ Statistics.

| Sample | Input Read Pairs | Both Surviving | Forward Only Surviving | Reverse Only Surviving | Dropped |
| --- | --- | --- | --- | --- | --- |
| Susceptible Control | 16933056 | 15309759<br>(90.41%) | 1297266<br>(7.66%) | 117401<br>(0.69%) | 208630<br>(1.23%) |
| Susceptible Infected | 17224957 | 15436022<br>(89.61%) | 1492707<br>(8.67%) | 111838<br>(0.65%) | 184390<br>(1.07%) |
| Susceptible treated | 20157934 | 18352456<br>(91.04%) | 1434433<br>(7.12%) | 146912<br>(0.73%) | 224133<br>(1.11%) |
| Resistant Control | 17666231 | 15819666<br>(89.55%) | 1513153<br>(8.57%) | 115125<br>(0.65%) | 218287<br>(1.24%) |
| **Resistant Infected** | 17678885 | 15639086<br>(88.46%) | 1707454<br>(9.66%) | 112901<br>(0.64%) | 219444<br>(1.24%) |
| **Resistant treated** | 23959344 | 21517460<br>(89.81%) | 1941133<br>(8.10%) | 146443<br>(0.61%) | 354308<br>(1.48%) |

**Table 3:**
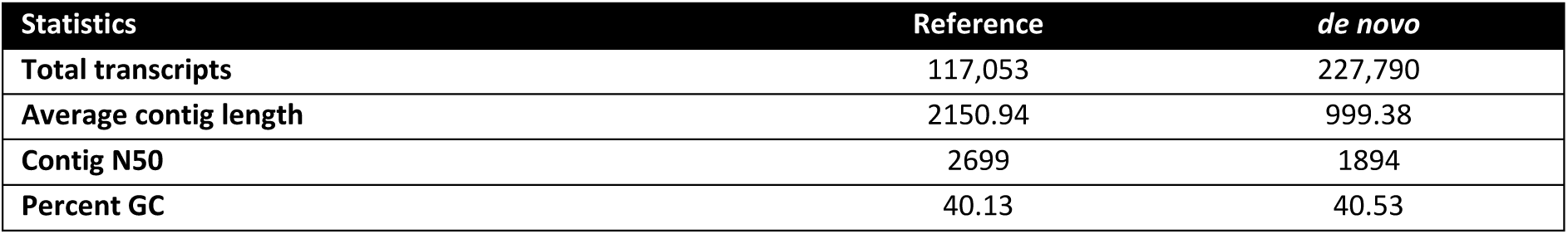
Statistical comparison of reference-based and *de novo* transcriptomic assembly of *Glycine max*.

| Statistics | Reference | *de novo* |
| --- | --- | --- |
| Total transcripts | 117,053 | 227,790 |
| Average contig length | 2150.94 | 999.38 |
| Contig N50 | 2699 | 1894 |
| Percent GC | 40.13 | 40.53 |

#### 2.6.1. Differentially expressed genes (DEGs) identification

In this study, significant DEGs with defined parameters (FDR and P-value <0.05 and log2fold change > 2) were identified in nine different comparison sets as presented in figure 5. A total of 2079 unique (3106 total in all conditions) DEGs were obtained which were distributed in nine comparison sets. The number of DEGs in each of the comparison sets are 354, 21, 318, 929, 412, 206, 41, 519 and 306 in susceptible-control vs resistant-control, susceptible-control vs susceptible-infected, susceptible-control vs susceptible-treated, susceptible-infected vs resistant-infected, susceptible-treated vs resistant-treated, resistant-control vs resistant-infected, resistant-control vs resistant-treated, resistant-infected vs resistant-treated and susceptible-infected vs susceptible-treated respectively (Figure 5, Supplementary Sheet 1). Heatmap showing the hierarchically clustered Spearman correlation matrix resulting from comparing the transcript expression values (TMM-normalized FPKM) for each sample is represented in Figure 6. Hierarchical clustering of the DEGs as well as samples is shown as a heatmap where clustering is based on transcripts abundance per sample (Supplementary Figure 1). Supplementary Figure 2 provides MA and volcano graphs of individual comparisons. The shared and unique differentially expressed genes are represented in the form of bar diagram in Figure 7. Highest number of transcripts (531) were differentially expressed in susceptible-infected vs resistant-infected indicated that the resistant variety has expressed resistant genes against the infection. The next highest number was observed in susceptible-treated vs resistant-treated (215) again indicated the presence of treatment sensitive genes. A total of 154, 5, 79, 531, 215, 30, 11, 207 and 85 DEGS were unique to the comparison sets susceptible-control vs resistant-control, susceptible-control vs susceptible-infected, susceptible-control vs susceptible-treated, susceptible-infected vs resistant-infected, susceptible-treated vs. resistant-treated, resistant-control vs resistant-infected, resistant-control vs resistant-treated, resistant-infected vs resistant-treated and susceptible-infected vs susceptible-treated respectively. 531 genes were found common in any 2 of the comparison, 203 genes where shared between 3 sets, 24 genes were found to be shared between four sets, 3 genes were shared between 5 sets while only one gene was shared between 7 sets. (Figure 7).

**Figure 5.**
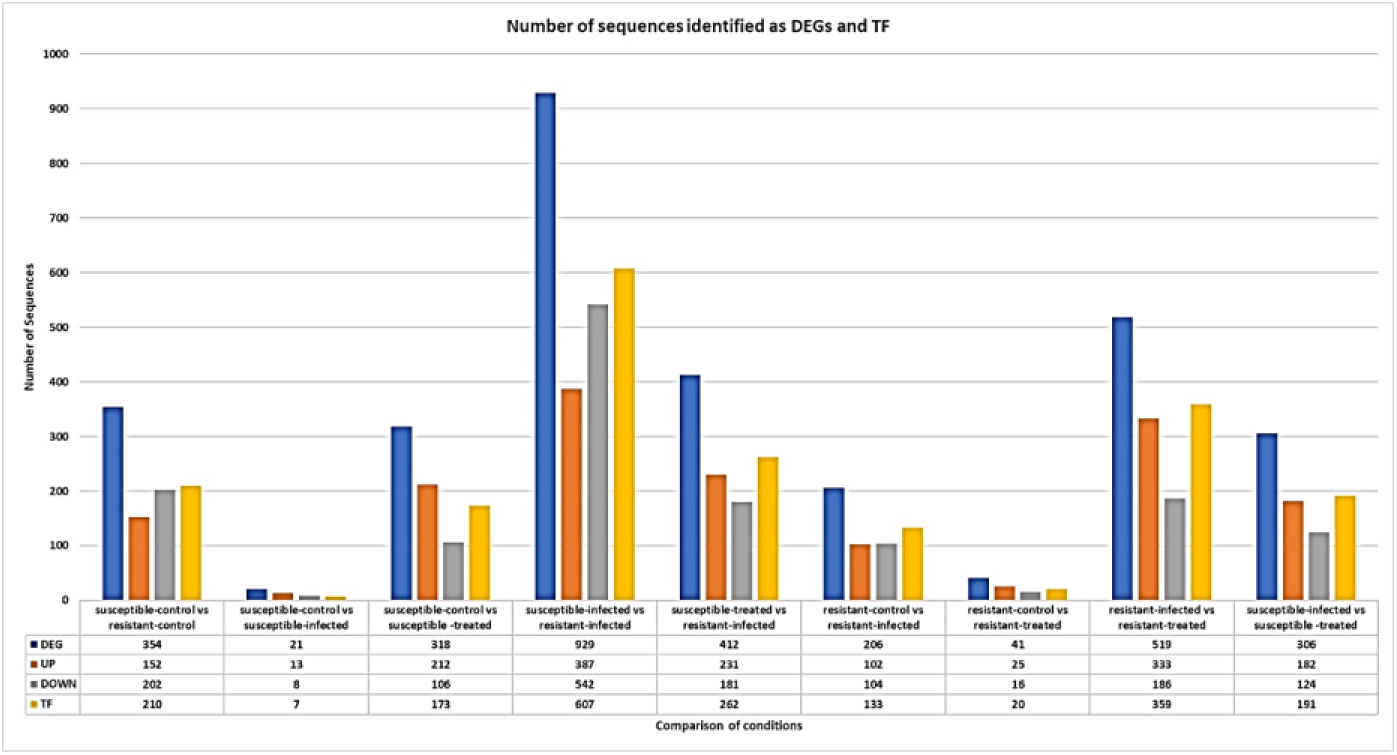
Number of significant differentially expressed genes and Transcription Factors in various comparison sets.

**Figure 6.**
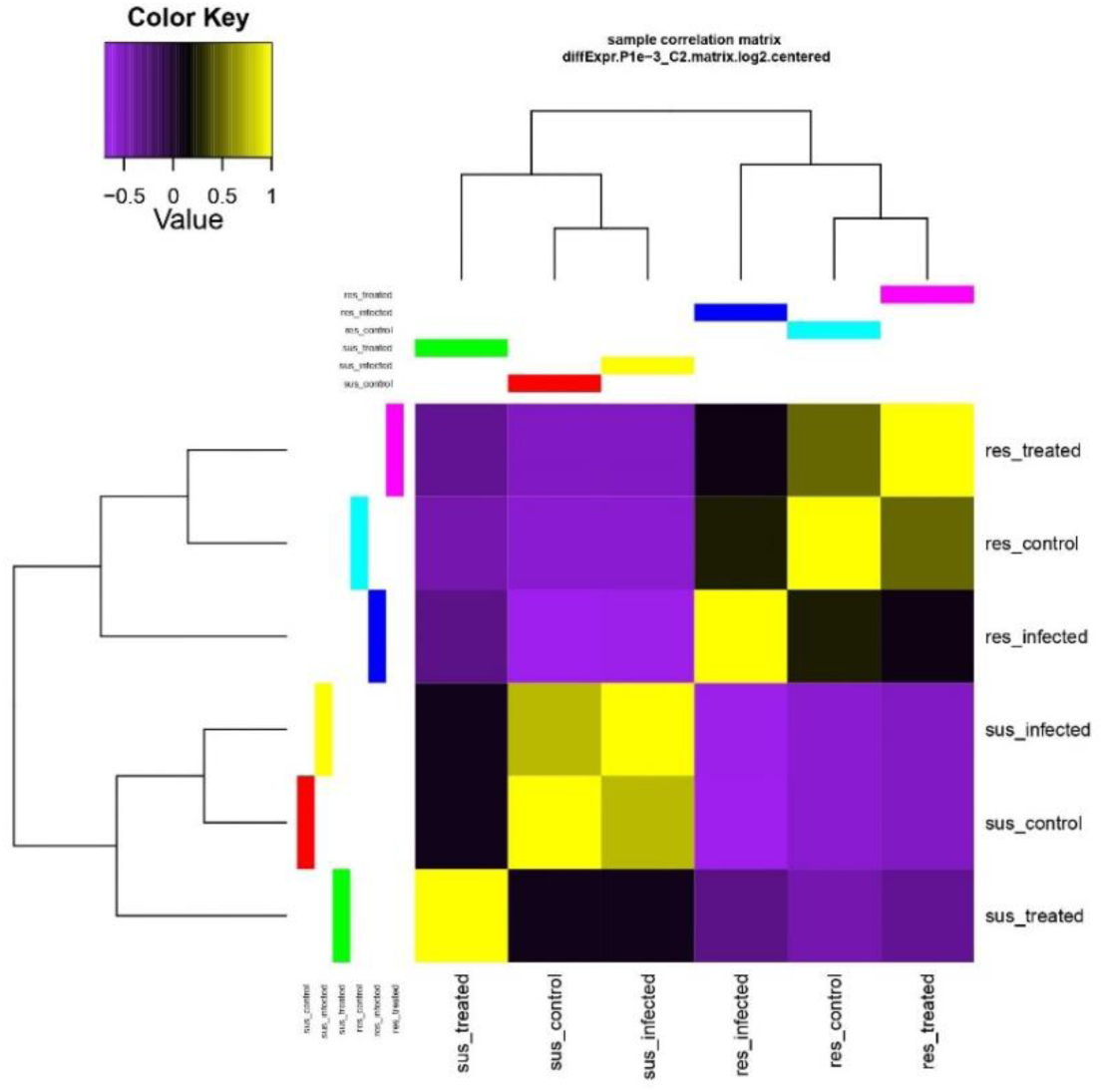
Heatmap showing the hierarchically clustered Spearman correlation matrix resulting from comparing the transcript expression values for each of the 6 samples.

**Figure 7.**
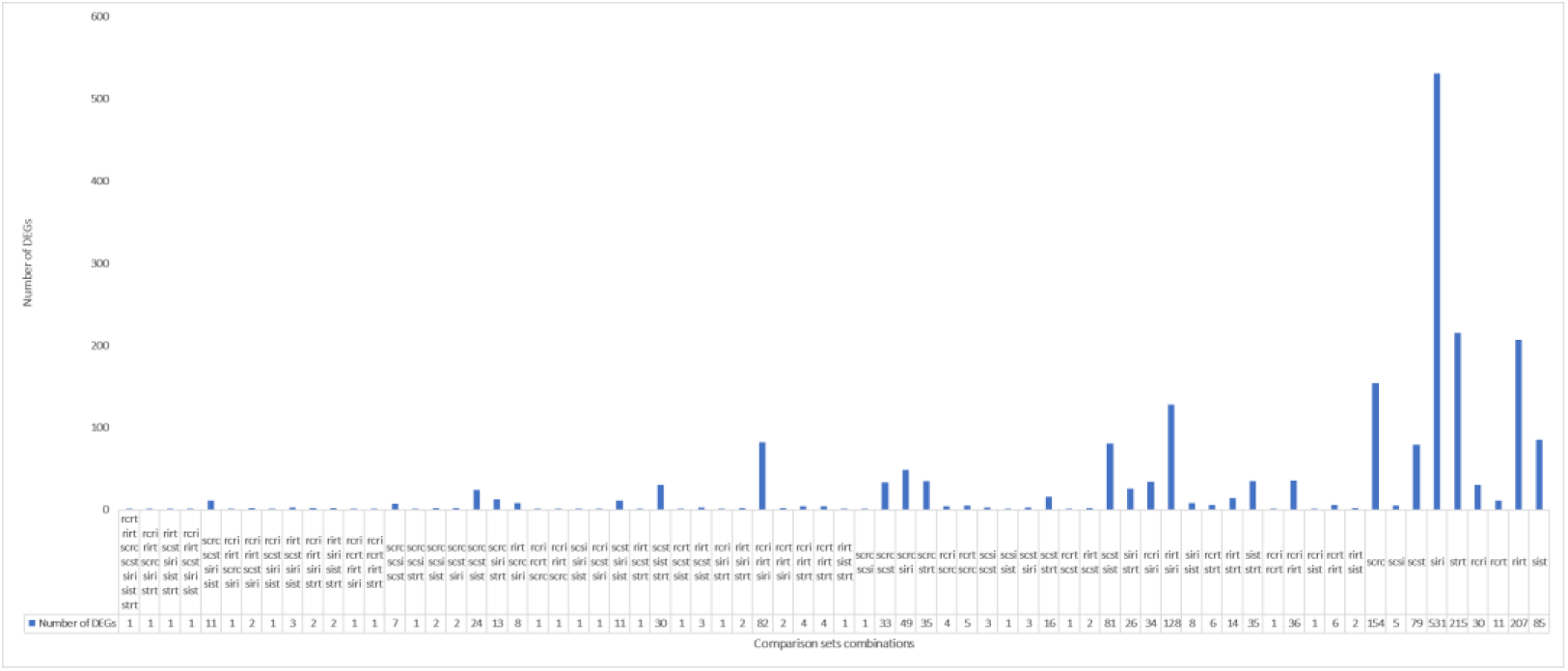
A bar diagram showing the DEGs in each of the nine comparison sets. Some DEGs are shared between different comparison sets

#### 2.6.2. Defense related genes

In order to establish a baseline, gene expression profiles from samples that were both healthy and infected along with 1.7mM Si treatment were used. If the gene’s expression differed by two folds (or more) in infected samples compared to control samples (p 0.001), the gene was considered to be a DEG (Table 4-6). The expression data of DEGs LRR receptor-like serine/threonine-protein kinase, NRT1/PTR FAMILY protein, ethylene-responsive transcription factor ABR1, cinnamyl alcohol dehydrogenase were noticeably elevated in resistant infected plants where they help in plant immunity boost up by recognizing pathogenic invaders. While alpha-amylase inhibitor, lipid transport, and seed storage family protein precursor, protein DETOXIFICATION 40 isoform-A, BURP domain protein RD22 genes are downregulated. The genes ORF16-lacZ fusion protein, putative oRF58e, and putative membrane protein are increased in the resistant treated plants, while inositol-3-phosphate synthase, NAC domain protein is downregulated shows the effectiveness of Si application. Senescence-associated protein, Thiol protease aleurain-like isoform A, multiple organellar RNA editing factor 9-like, and zinc finger protein ZAT10-like were dramatically elevated genes in the susceptible infected plants. The genes Protein SRC1-like and Early Nodulin-Like Protein 1 isoform B are upregulated in the susceptible treated plants, whereas Proline-Rich Protein 2 isoform C and B, Gibberellin-Regulated Protein 6 (like), Repetitive Proline-Rich Cell Wall Protein 1, and plant senescence-associated protein, putative are downregulated genes. Many genes in susceptible treated plants were associated with hypothetical proteins for which comprehensive annotations were not yet available. The gene carbonic anhydrase, chloroplastic isoform 1 was significantly increased in susceptible plants compared to resistant plants, but it was significantly downregulated in resistant plants. A gene that encodes RNA-binding protein 12 activates stress-responsive mRNAs was upregulated in susceptible plants and downregulated in resistant ones there must be linking to disease development. In susceptible and resistant plants, respectively, there was upregulation and downregulation of the auxin-binding protein ABP19a-like gene. Our results indicate that up-regulation of defense-related genes and enzymes enhanced resistance to *Macrophomina phaseolina* infection without having a negative impact on soybean plant growth and development. The upregulation of pathogenesis related protein (PRP) genes in resistant plants strongly indicates the involvement of a resistant gene within this domain of PR proteins. The activation of programmed cell death in susceptible plants following silicon treatment suggests that certain genes are triggered to halt the spread of fungal infections by initiating the programmed cell death process. Silicon act as a priming agent, preparing the plant to respond more effectively to subsequent pathogen attacks.

**Table 4.** Differentially expressed transcripts in resistant infected and susceptible infected plants.

| Sr.no. | Gene ID | Description | Resistant infected | Susceptible infected |
| --- | --- | --- | --- | --- |
| 1 | TRINITY_DN152677_c0_g3_i2 | XP_003545772.1pathogenesis-related protein 1 | Upregulated | Downregulated |
| 2 | TRINITY_DN157192_c1_g2_i4 | NP_001236055.1stress-induced protein H4 | Upregulated | Downregulated |
| 3 | TRINITY_DN163707_c2_g2_i2 | NP_001235512.1disease resistance protein | Upregulated | Downregulated |
| 4 | TRINITY_DN157192_c1_g1_i4 | XP_003543663.1 pleiotropic drug resistance protein 1 | Upregulated | Downregulated |
| 5 | TRINITY_DN157572_c5_g4_i2 | NP_001235031.1 BSP domain-containing protein precursor | Upregulated | Downregulated |
| 6 | TRINITY_DN163589_c2_g3_i2 | XP_003551991.1protein NRT1/ PTR FAMILY 2.13 | Upregulated | Downregulated |
| 7 | TRINITY_DN163688_c2_g6_i1 | NP_001238536.1 receptor-like kinase | Upregulated | Downregulated |
| 8 | TRINITY_DN160984_c2_g4_i1 | KHN03328.1 Major allergen Pru av 1 | Upregulated | Downregulated |
| 9 | TRINITY_DN159419_c0_g1_i5 | NP_001236953.2 dehydration responsive element-binding protein 3 | Upregulated | Downregulated |
| 10 | TRINITY_DN157005_c0_g2_i1 | NP_001238561.1chitinase class I precursor | Upregulated | Downregulated |
| 11 | TRINITY_DN154304_c2_g5_i2 | XP_003528760.1 anthranilate N-methyltransferase | Upregulated | Downregulated |
| 12 | TRINITY_DN157352_c1_g2_i5 | XP_003537598.1 isoflavone reductase | Upregulated | Downregulated |
| 13 | TRINITY_DN155040_c0_g1_i2 | NP_001237509.1serine hydroxymethyltransferase 5 | Upregulated | Downregulated |
| 14 | TRINITY_DN155388_c1_g1_i1 | XP_028238106.1probable protein phosphatase 2C 58 | Upregulated | Downregulated |

**Table 5.** Differentially expressed transcripts in resistant infected and resistant treated plants.

| Sr.no. | Gene ID | Description | Resistant infected | Resistant treated |
| --- | --- | --- | --- | --- |
| 1 | TRINITY_DN153404_c1_g1_i4 | XP_003551914.1trifunctional UDP-glucose 4,6-dehydratase/UDP-4-keto-6-deoxy-D-glucose 3,5-epimerase/UDP-4-keto-L-rhamnose-reductase RHM1 | Upregulated | Downregulated |
| 2 | TRINITY_DN148173_c0_g1_i1 | XP_028205173.1RNA-binding protein 12-like | Upregulated | Downregulated |
| 3 | TRINITY_DN154478_c0_g2_i4 | NP_001238515.22-hydroxyisoflavanone synthase precursor | Upregulated | Downregulated |
| 4 | TRINITY_DN157192_c1_g2_i2 | NP_001236038.1stress-induced protein SAM22 | Upregulated | Downregulated |
| 5 | TRINITY_DN37574_c0_g1_i1 | XP_003543246.1defensin-like protein | Upregulated | Downregulated |
| 6 | TRINITY_DN155588_c1_g7_i3 | NP_001237911.1SRPBCC ligand-binding domain-containing protein | Upregulated | Downregulated |
| 7 | TRINITY_DN155028_c2_g4_i2 | P10742.2RecName: Full=Stem 31 kDa glycoprotein; AltName: Full=Vegetative storage protein VSP25; Flags: Precursor | Downregulated | Upregulated |
| 8 | TRINITY_DN161100_c2_g6_i1 | NP_001238401.1alpha-amylase inhibitor/lipid transfer/seed storage family protein precursor | Downregulated | Upregulated |
| 9 | TRINITY_DN153400_c0_g1_i6 | XP_003522684.1BURPdomain protein RD22 | Downregulated | Upregulated |
| 10 | TRINITY_DN162634_c2_g1_i9 | XP_003531361.1inositol-3-phosphate synthase | Downregulated | Upregulated |
| 11 | TRINITY_DN162634_c2_g1_i7 | XP_028233150.1inositol-3-phosphate synthase | Downregulated | Upregulated |
| 12 | TRINITY_DN155028_c2_g4_i2 | P10742.2 Stem 31 kDa glycoprotein; | Downregulated | Upregulated |
| 13 | TRINITY_DN160334_c2_g2_i1 | NP_001344954.1germin-like protein precursor | Downregulated | Upregulated |
| 14 | TRINITY_DN151581_c0_g1_i1 | XP_028233150.1inositol-3-phosphate synthase | Downregulated | Upregulated |

**Table 6.**
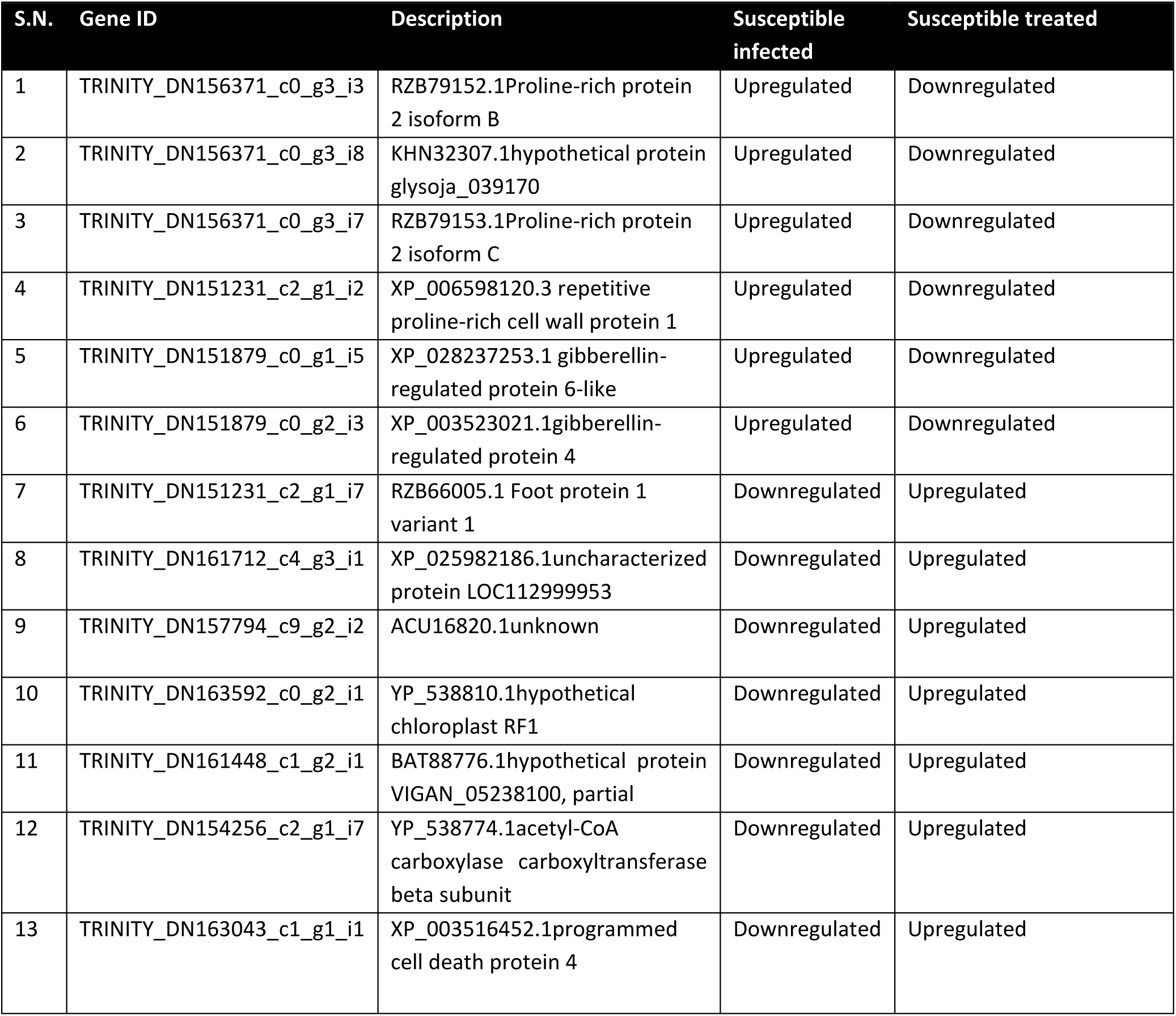
Differentially expressed transcripts in resistant infected and susceptible treated plants.

| S.N. | Gene ID | Description | Susceptible infected | Susceptible treated |
| --- | --- | --- | --- | --- |
| 1 | TRINITY_DN156371_c0_g3_i3 | RZB79152.1Proline-rich protein 2 isoform B | Upregulated | Downregulated |
| 2 | TRINITY_DN156371_c0_g3_i8 | KHN32307.1hypothetical protein glysoja_039170 | Upregulated | Downregulated |
| 3 | TRINITY_DN156371_c0_g3_i7 | RZB79153.1Proline-rich protein 2 isoform C | Upregulated | Downregulated |
| 4 | TRINITY_DN151231_c2_g1_i2 | XP_006598120.3 repetitive proline-rich cell wall protein 1 | Upregulated | Downregulated |
| 5 | TRINITY_DN151879_c0_g1_i5 | XP_028237253.1 gibberellin-regulated protein 6-like | Upregulated | Downregulated |
| 6 | TRINITY_DN151879_c0_g2_i3 | XP_003523021.1gibberellin-regulated protein 4 | Upregulated | Downregulated |
| 7 | TRINITY_DN151231_c2_g1_i7 | RZB66005.1 Foot protein 1 variant 1 | Downregulated | Upregulated |
| 8 | TRINITY_DN161712_c4_g3_i1 | XP_025982186.1uncharacterized protein LOC112999953 | Downregulated | Upregulated |
| 9 | TRINITY_DN157794_c9_g2_i2 | ACU16820.1unknown | Downregulated | Upregulated |
| 10 | TRINITY_DN163592_c0_g2_i1 | YP_538810.1hypothetical chloroplast RF1 | Downregulated | Upregulated |
| 11 | TRINITY_DN161448_c1_g2_i1 | BAT88776.1hypothetical protein VIGAN_05238100, partial | Downregulated | Upregulated |
| 12 | TRINITY_DN154256_c2_g1_i7 | YP_538774.1acetyl-CoA carboxylase carboxyltransferase beta subunit | Downregulated | Upregulated |
| 13 | TRINITY_DN163043_c1_g1_i1 | XP_003516452.1programmed cell death protein 4 | Downregulated | Upregulated |

#### 2.6.3. Homology search, annotation, functional characterization and transcription factor identification

All the sequences in the above mentioned nine comparison sets were subjected to homology search using Blast2GO. Highest similarity was identified against Glycine soja (>1250 sequences) followed by Glycine max (>1150 sequences) (Supplementary Figure 3) which proves their phylogenetic relatedness. Gene ontology (GO) classification of all the DEGs was also carried out. For the nine sets that were compared i.e. (1) susceptible-control vs resistant-control (2) susceptible-control vs susceptible-infected (3) susceptible-control vs susceptible-treated (4) susceptible-infected vs resistant-infected (5) susceptible-treated vs resistant-treated (6) resistant-control vs resistant-infected (7) resistant-control vs resistant-treated (8) resistant-infected vs resistant-treated and (9) susceptible-infected vs susceptible-treated, all the GO terms were identified and respectively are as follows (Total:CC-MF-BP): (1) 382:61-163-158 (2) 54:14-22-18 (3) 390:65-158-167 (4) 635:78-279-278 (5) 584:94-210-280 (6) 201:30-97-74 (7) 70:11-34-25 (8) 422:54-184-184 (9) 426:64-176-186 (Supplementary Sheet 2). It was observed that most of the transcripts annotated under cellular components belong to the membrane ontology with ‘integral component of membrane’ (GO:0016021) and membrane (GO:0016020) being the top two. The highest molecular function assigned was from transferase activity (GO:0016740). ‘Regulation of transcription’ (GO:0006355) was observed to be the highest in biological processes followed by ‘transmembrane transport’ (GO:0055085) (Figure 8).

**Figure 8.**
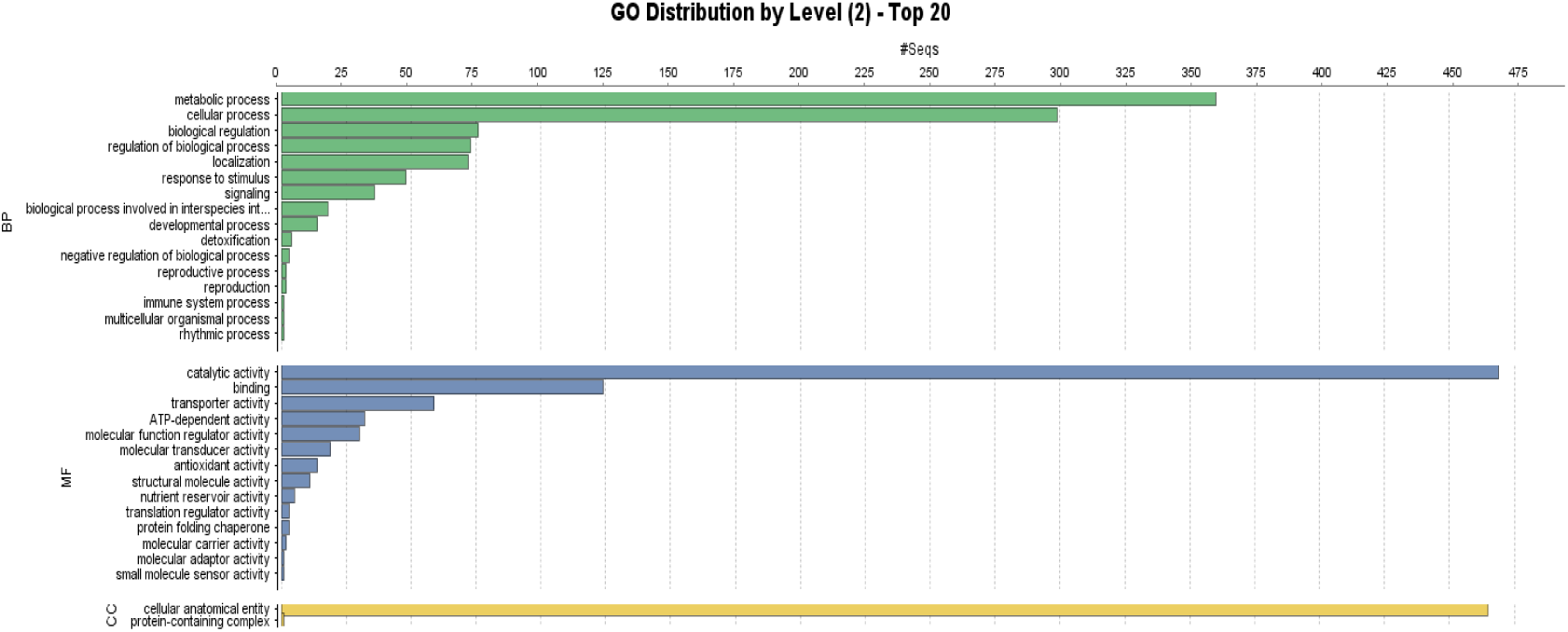
GO distribution of DEGs.

GO enrichment analysis was performed for all the upregulated and downregulated DEGs separately. In case of upregulated genes and it was found that in GO category Biological Processes ‘lipid metabolic process’ (GO:0005200) has the highest percentage of enriched GO terms (5.12). In case of Cellular Components ‘extracellular regions’ (GO:0005576) has the highest percentage of 5.71 followed by ‘vacuole’ (GO:0004512) (2.36). In Molecular functions ‘structural constituent of cytoskeleton’ (GO:0006021) have the highest sequence percentage (1.08). All the details have been shown in Figure 9 and Supplementary sheet 3. In case of downregulated genes Biological Processes GO term ‘transmembrane transport’ (GO:0031347) having the highest percentage of enriched genes (9.38), Cellular Components has only one GO term ‘plasma membrane’ with 10.68 % of enriched genes while in Molecular functions ‘catalytic activity’ has a high percentage of enriched sequences (13.53) (Figure 10, Supplementary sheet 3).

**Figure 9.**
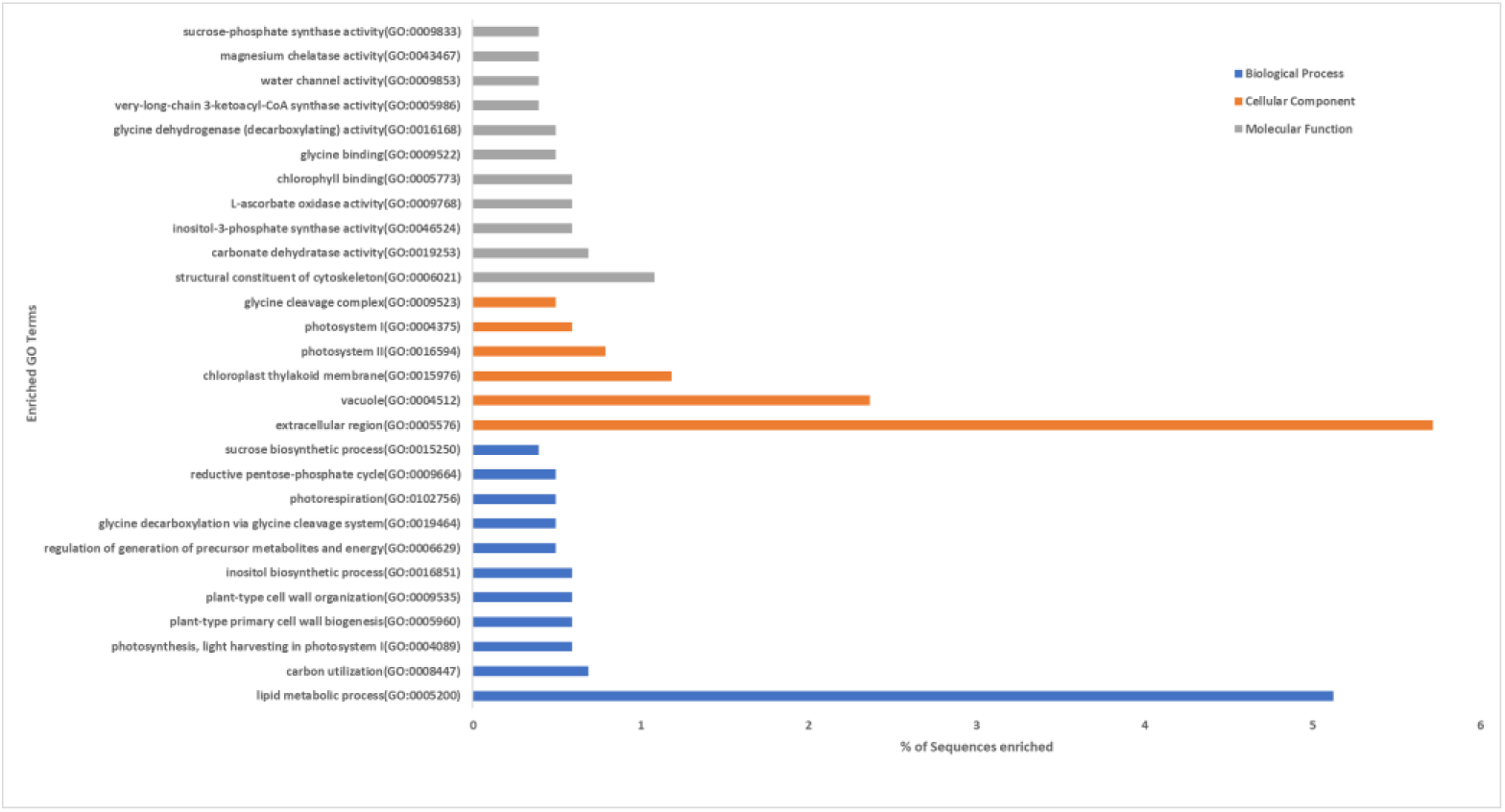
GO enrichment of upregulated DEGs by Fisher’s Exact Test.

**Figure 10.**
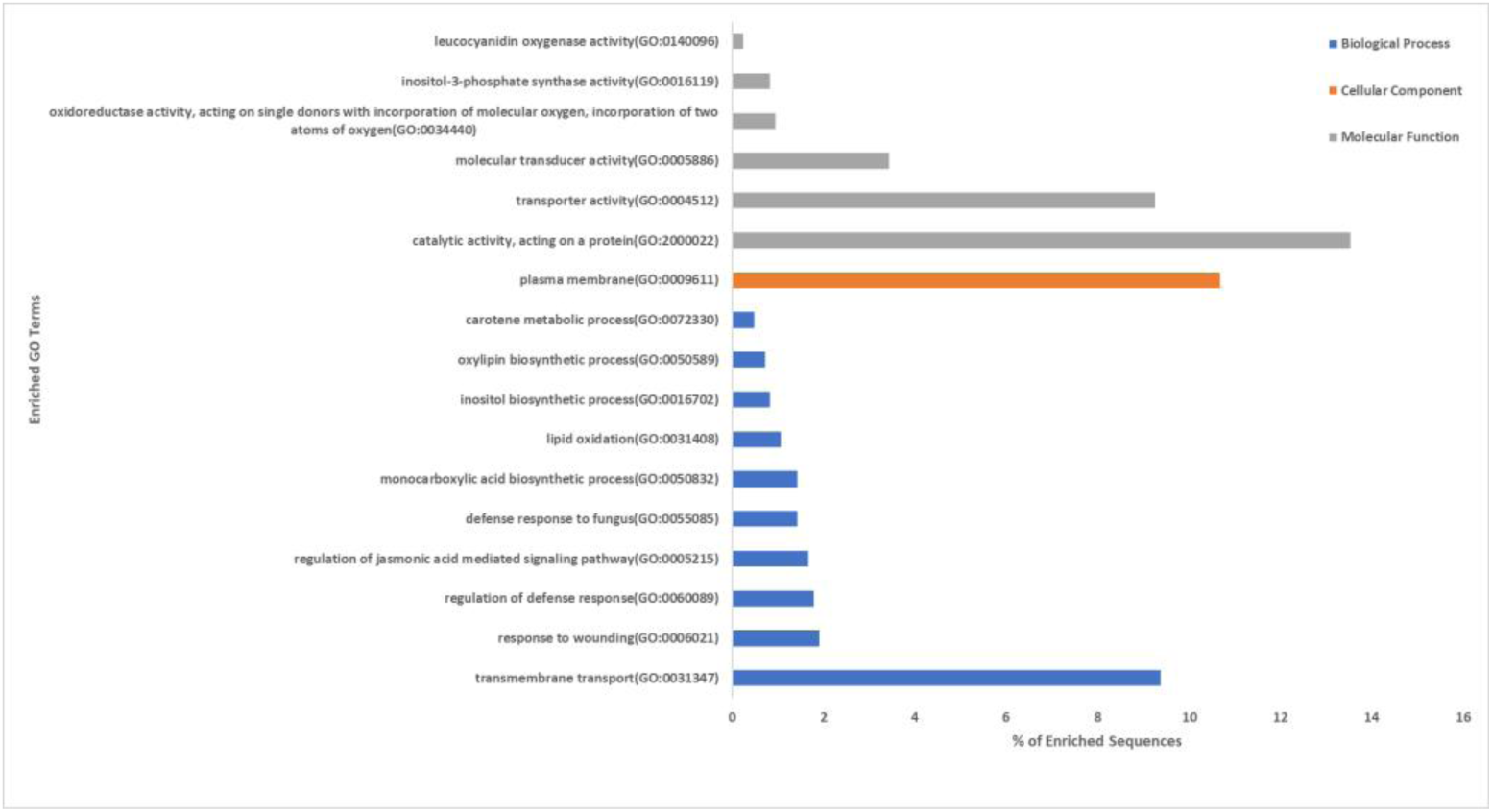
GO enrichment of downregulated DEGs by Fisher’s Exact Test.

Kyoto Encyclopedia of Genes and Genomes (KEGG) pathway analysis revealed that from a total of 354, 21, 318, 929, 412, 206, 41, 519 and 306 DEGs obtained from nine comparisons 22, 0, 21, 71, 23, 16, 2, 32 and 15 transcripts were assigned to one or more pathways respectively. It was observed that a high number of sequences were present in pathways related to sugar metabolism like glycolysis/gluconeogenesis (map00010) and starch and sucrose metabolism (map00500) (Supplementary Sheet 4). Figure 11 shows the top 30 KEGG pathways identified in the DEGs. KEGG enrichment analysis revealed that KEGG pathway ‘Biosynthesis of ansamycins (map01051) is highly enriched (rich factor = 0.33) followed by ‘Flavone and flavonol biosynthesis (map00944) (rich factor = 0.25) (Figure 12). We identified a total of 210, 7, 173, 607, 262, 133, 20, 359 and 191 transcriptions factors (TFs) in DEGs obtained from nine sets of comparisons (Figure 5). The highest number of sequences has been reported with ERF (Ethylene responsive factor) transcription factor followed by helix loop helix family protein (bHLH). Other important TF identified are MYB, NAC and FAR1 family proteins (Supplementary Table 5).

**Figure 11.**
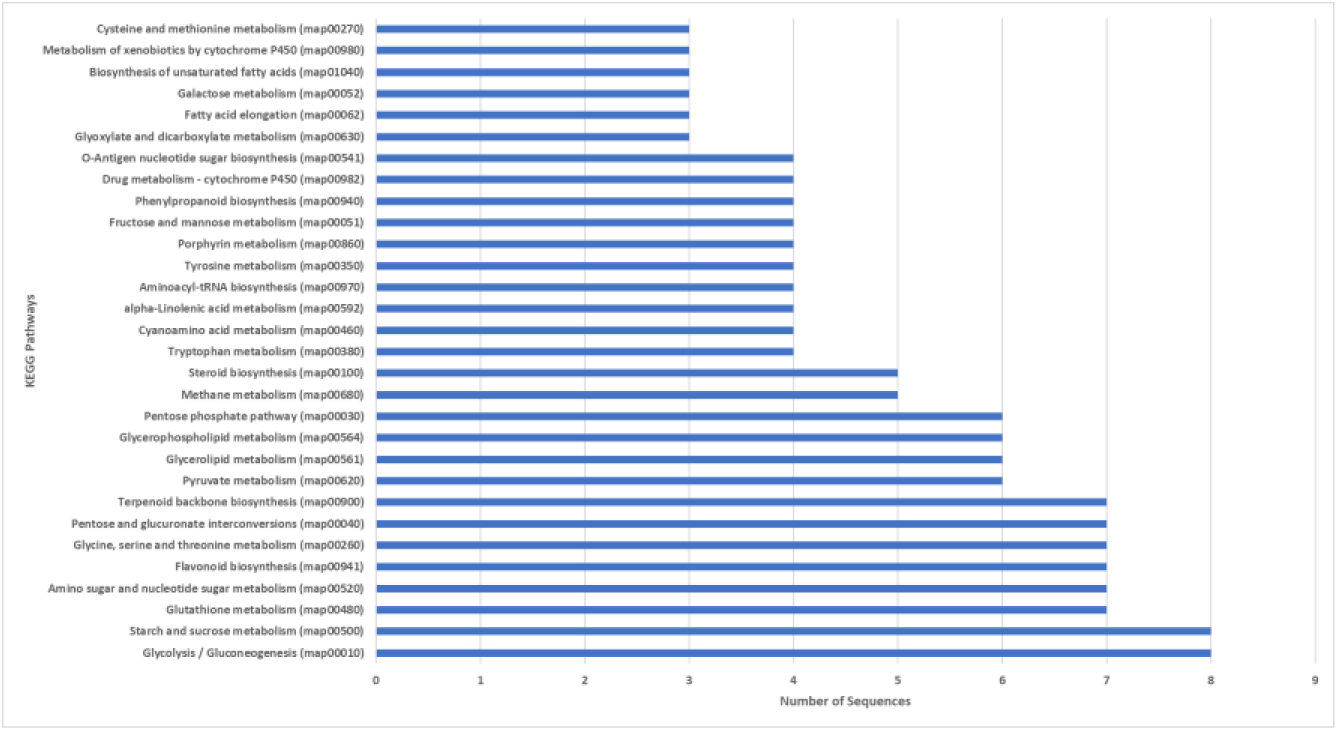
Top 30 KEGG pathways identified based on the number of sequences present in that pathway.

**Figure 12.**
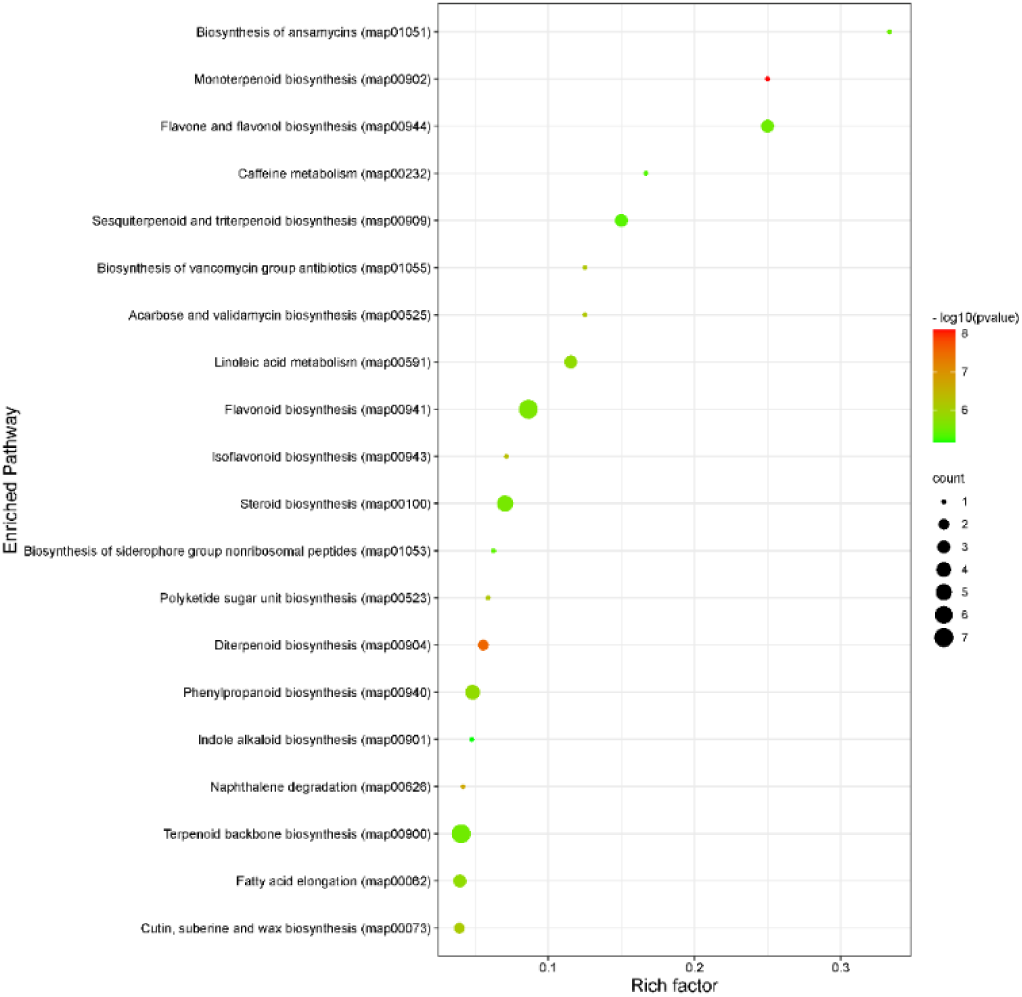
KEGG pathway enrichment[13].

#### 2.6.4. Variant identification

Based on the de novo assembled transcripts, a total of 1,11,447 putative SSRs were identified. The primers were designed for 1,10,999 SSRs while for 448 SSRs primers could not be designed. Out of the SSRs for which primers were designed mononucleotides were 65,742 (59.22%), dinucleotides were 18,471(16.64%), trinucleotide were 16,369(14.74%), tetranucleotide were 1185(1.06%), pentanucleotide were 287(0.25%), hexanucleotide were 193(0.17%) and compound SSR were 8752(7.88%). Supplementary Table 6 gives the details of all the identified SSR markers and their primers. SNPs and InDels were identified in the two contrasting varieties i.e., resistant and susceptible. A total of 518,501 raw variants were detected in the resistance variety (including SNP, MNP, Insertion, Deletions and complex variants) while in the susceptible variety 442,267 raw variants were detected. List of variants has been provided in Supplementary Table 7. A circus plot of all the variants is shown in Figure 13.

**Figure 13.**
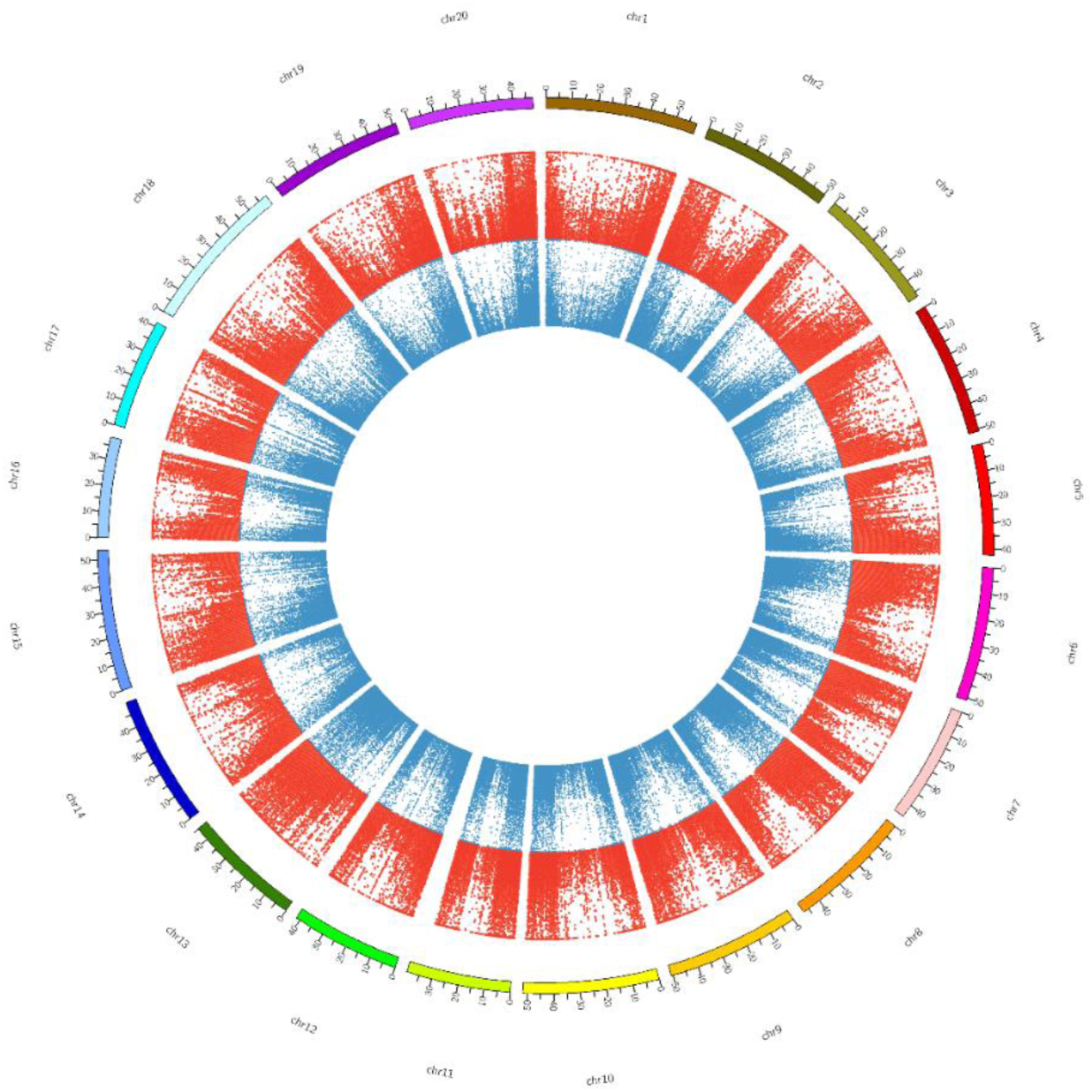
Circos plot of all the variants (ANP and INDEL) present in resistant (blue) and susceptible (red) variety.

#### 2.6.5. Gene regulatory networks

The list of top 50 genes (25 upregulated and 25 downregulated) from each of the nine sets were used for construction of gene regulatory networks (GRNs) as provided in Supplementary Table 8. In the gene regulatory network figures, genes represented in red are upregulated while those in blue are downregulated. Genes in yellow have been identified as potential hub genes in the network. All the gene networks have been shown in Figure 14(a-g). Heatmaps were also generated for all the DEGs present in the nine GRNs (figure 15(a-i)). These heatmaps can pinpoint exclusive genes which show high fold change in a comparison.

**Figure 14.**
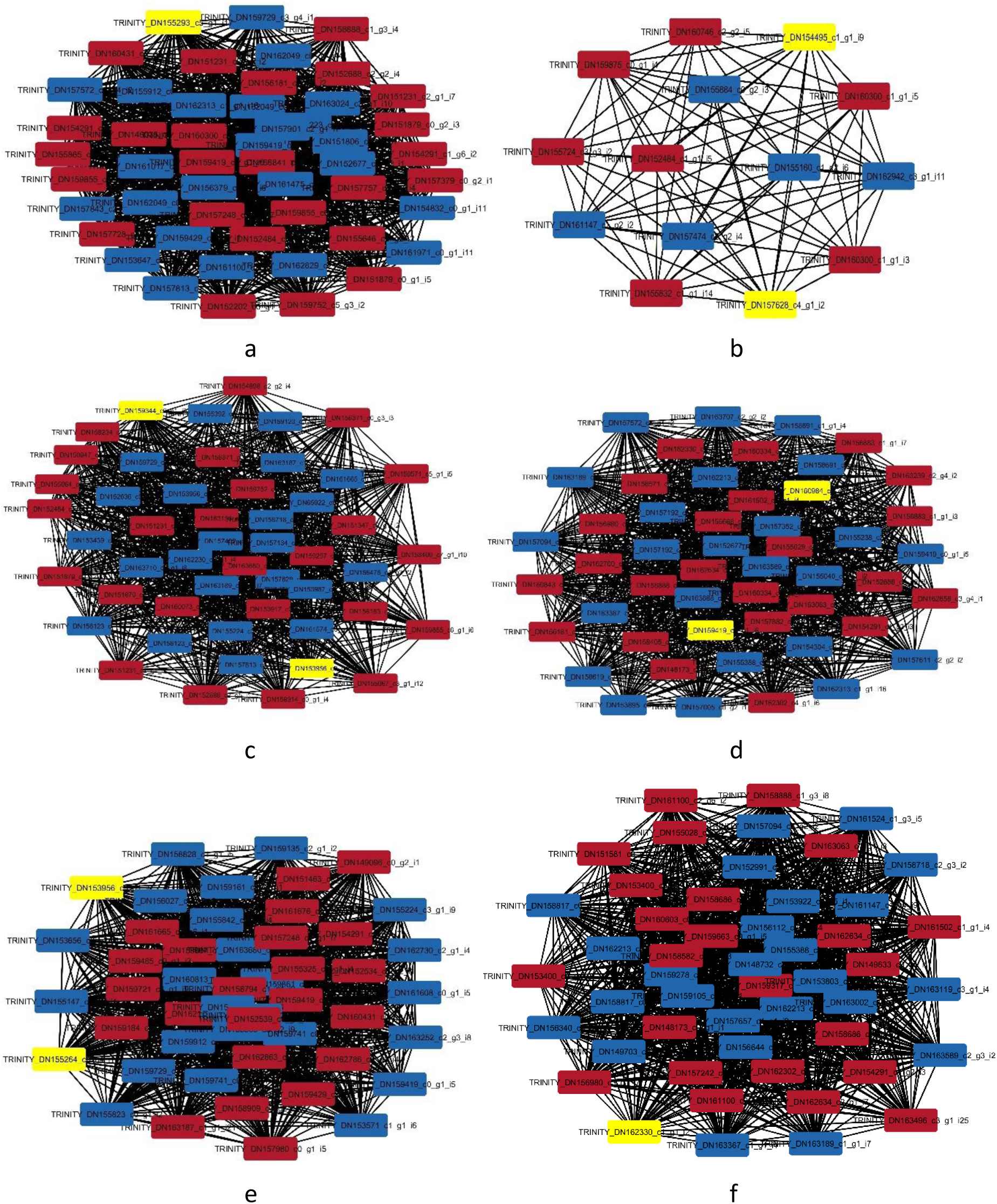
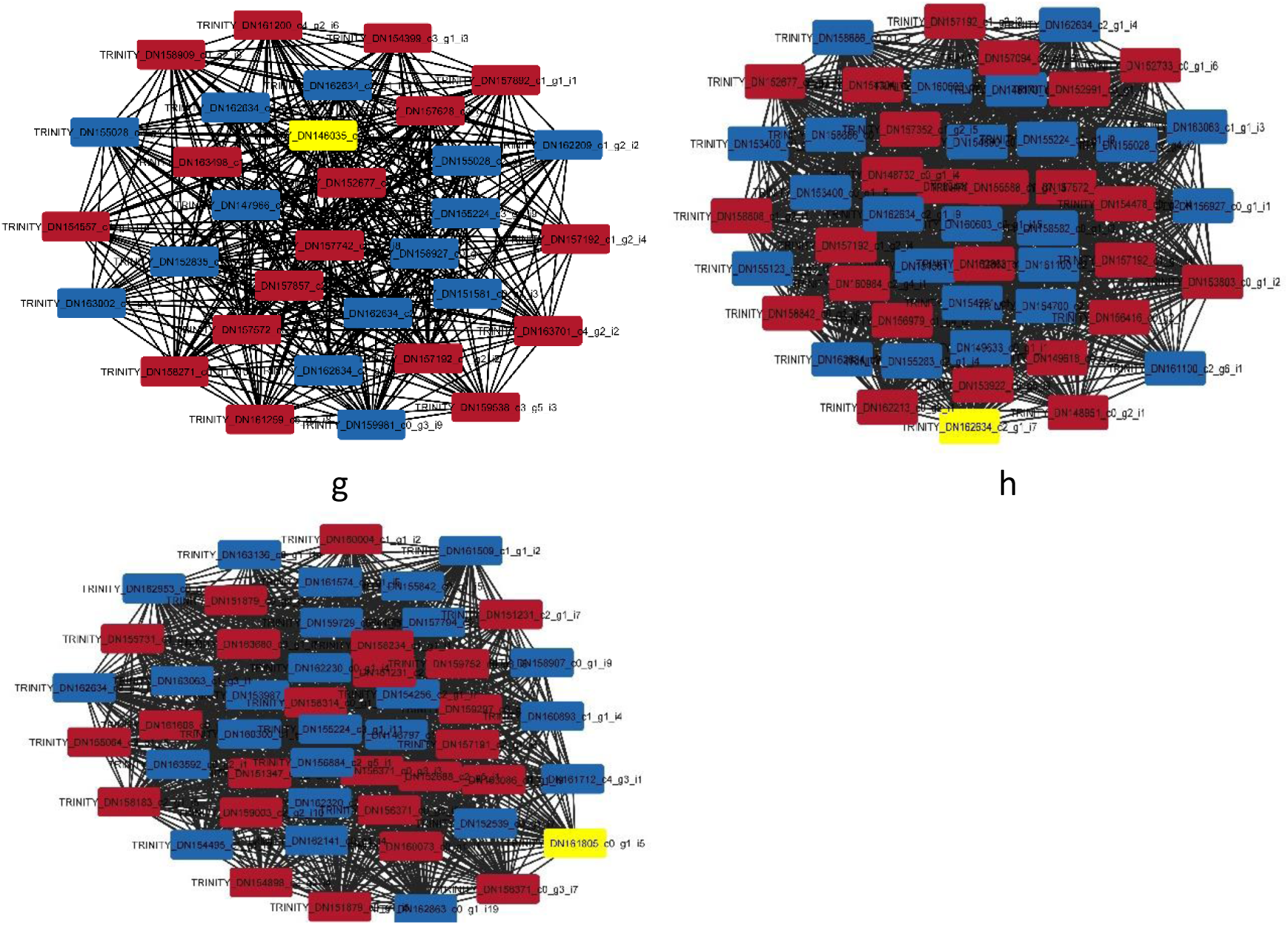
Gene regulatory networks as identified in nine comparison sets. (a) susceptible-control vs resistant-control (b) susceptible-control vs susceptible-infected (c) susceptible-control vs susceptible-treated (d) susceptible-infected vs resistant-infected (e) susceptible-treated vs resistant-treated (f) resistant-control vs resistant-infected (g) resistant-control vs resistant-treated (h) resistant-infected vs resistant-treated and (i) susceptible-infected vs susceptible-treated.

**Figure 15.**
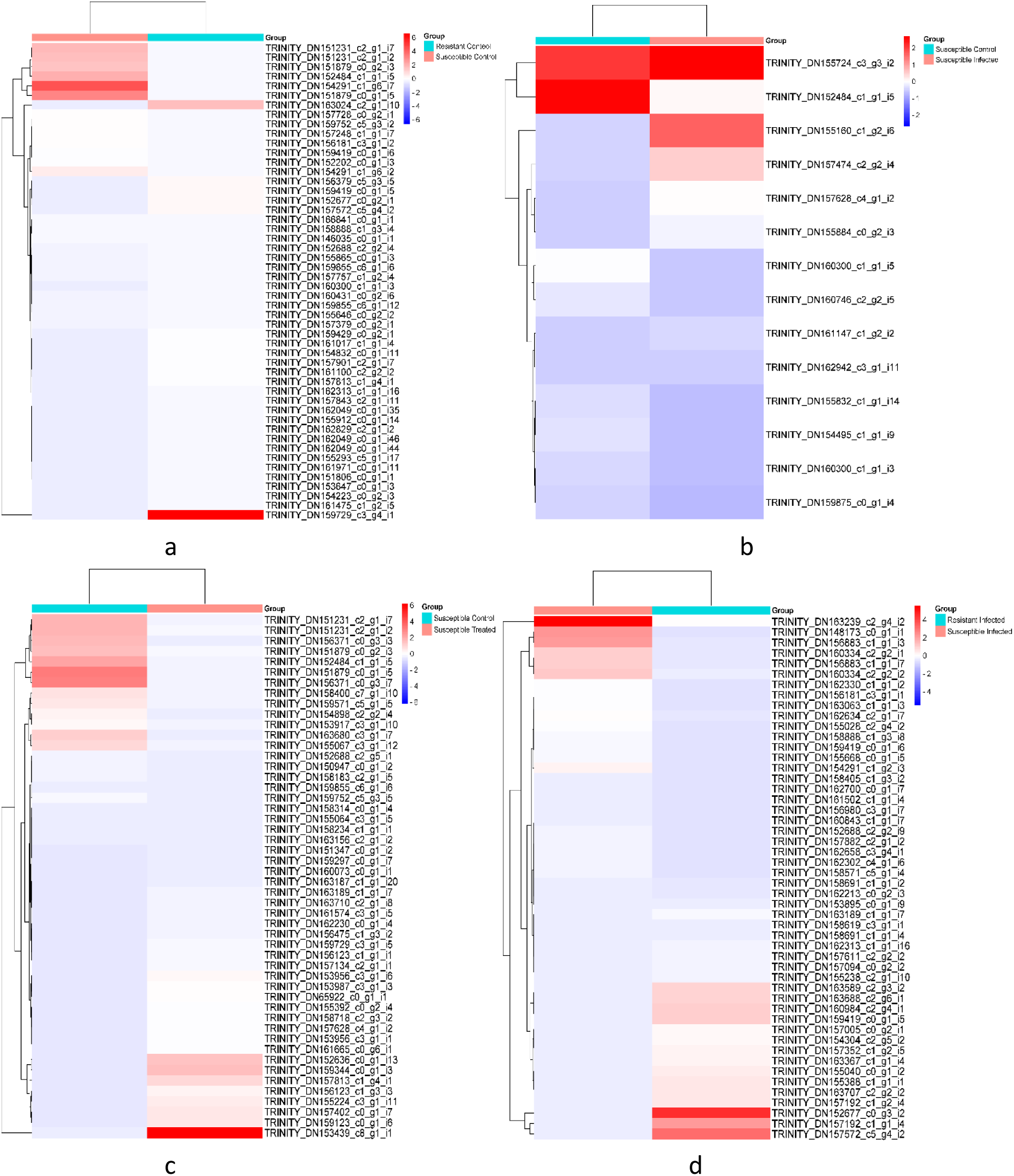
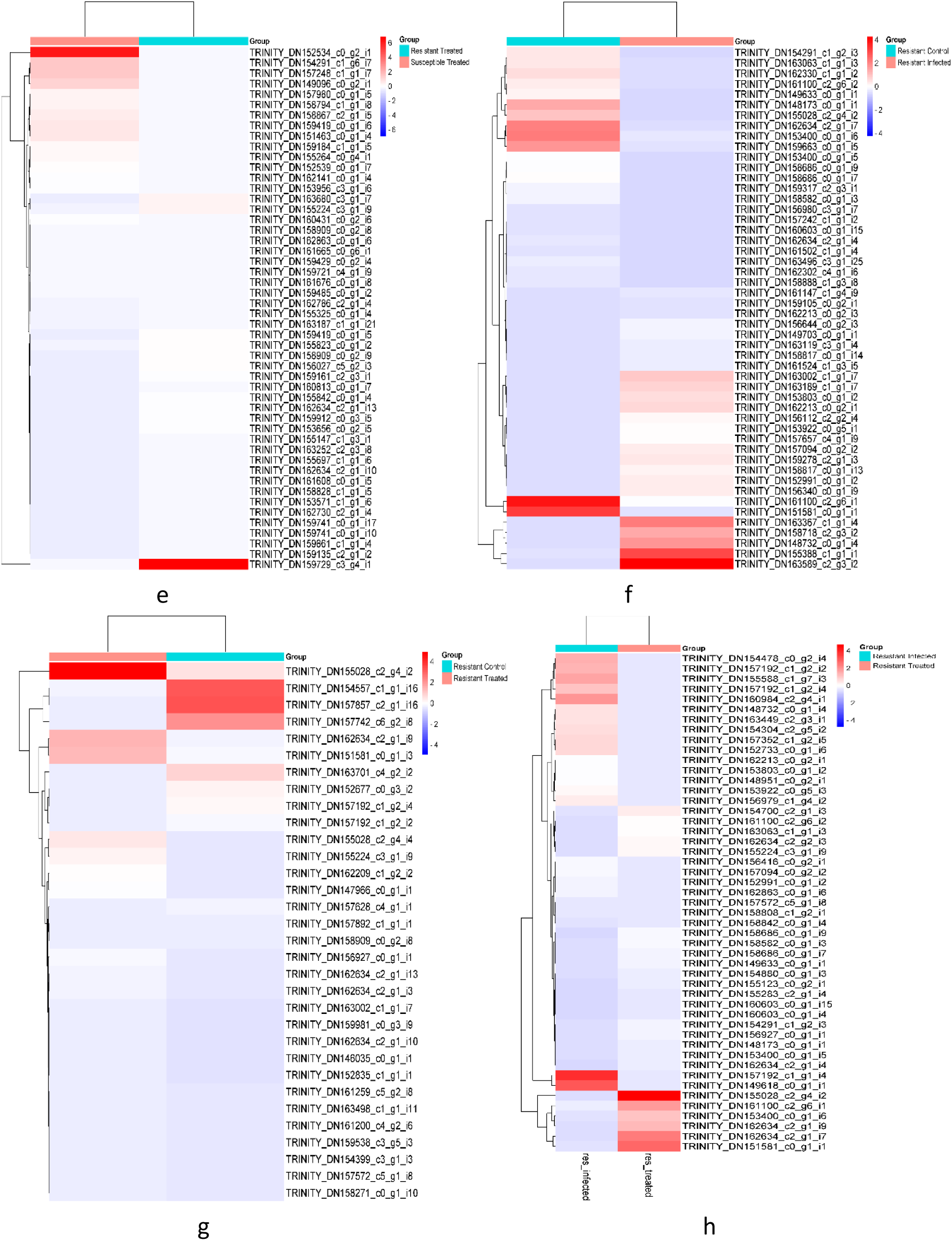
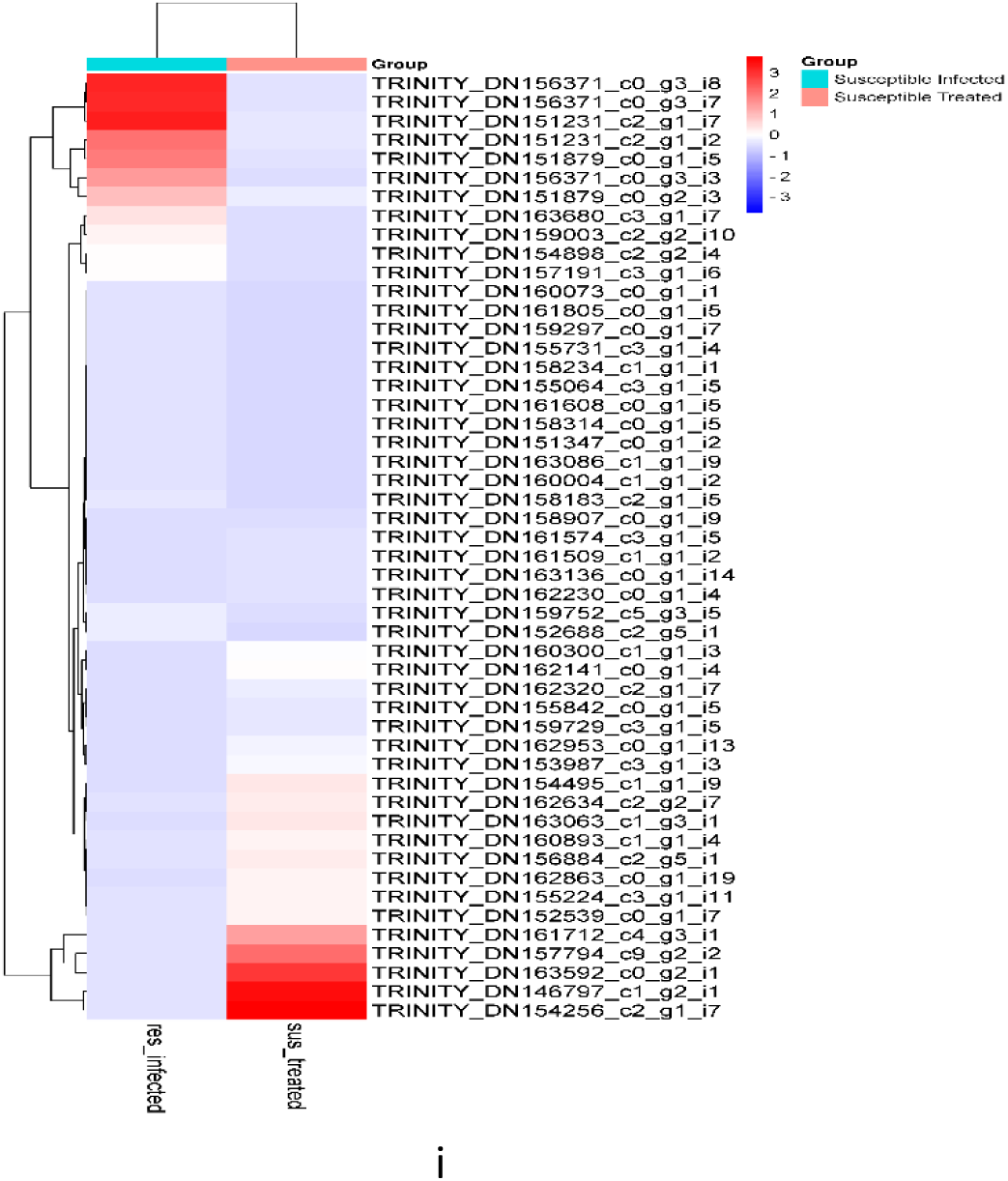
Heatmaps for DEGs present in Gene regulatory networks. (a) susceptible-control vs resistant-control (b) susceptible-control vs susceptible-infected (c) susceptible-control vs susceptible-treated (d) susceptible-infected vs resistant-infected (e) susceptible-treated vs resistant-treated (f) resistant-control vs resistant-infected (g) resistant-control vs resistant-treated (h) resistant-infected vs resistant-treated and (i) susceptible-infected vs susceptible-treated.

#### 2.6.6. Soybean Transcriptomics database

Data from the examination of transcriptomics were abundant. A website resource (STDbCr) has been created for soybean for the general public to utilize (http://backlin.cabgrid.res.in/stdbcr/). The actual transcripts that were created, DEGs that were found in different comparison sets, transcription factors, and biomarkers like SSR and SNP/InDels are all stored in this database. The data is presented in an interactive table, which enables the user to search or filter the data appropriately. Currently, there are 3,106 DEGs (spread over nine comparisons), 10,388 transcription factors, 110,999 SSR, 518,501 variations in the resistant variety, and 442,267 variants in the susceptible variety, in the database. Users may browse the data in this database since it is made to be simple to use. Links to the initial data delivered to NCBI are available in a download section. Additionally, contact information has been given. The RNAseq data for the study has been deposited in a repository with the BioProject title “The RNAseq Study on Charcoal Rot Fungal Disease in Soybean.” This project has been assigned the accession number PRJNA1026744 and the corresponding ID 1026744. BioSample: **SAMN37757654**: TAMS-38_S1: Control (Normal watering as per requirement) without charcoal rot infection):SRR26357052; **SAMN37757655**: TAMS-38_S2 (Charcoal rot infected soil (Normal watering as per requirement)):SRR26357051; **SAMN37757656**: TAMS-38_S3 (Charcoal rot infected soil along with 1.7mM K_2_SiO_3_ treatment (1.7mM K_2_SiO_3_ treatment by water drenching)):SRR26357050; **SAMN37757657**: AMS-MB-5-18_S4 (Control (Normal watering as per requirement) without charcoal rot infection):SRR26357049; **SAMN37757658**: AMS-MB-5-18_S5 (Charcoal rot infected soil (Normal watering as per requirement)): SRR26357048; **SAMN37757659**: AMS-MB-5-18_S6 (Charcoal rot infected soil along with 1.7mM K_2_SiO_3_ treatment (1.7mM K_2_SiO_3_ treatment by water drenching)): SRR26357047. The data can be access by following this link: (https://www.ncbi.nlm.nih.gov/bioproject/?term=PRJNA1026744)

#### 2.6.7. Validation of DEGs by qRT-PCR

Quantitative real-time PCR of 8 randomly chosen DEGs was carried out to verify genes retrieved from RNA-seq data. The gene-specific primers were designed and examined in samples of soybean that were susceptible, resistant, treated, and untreated (Table 7). The internal control Con-15 (a protein kinase related to CDPK) gene was utilized to normalize the amount of gene expression. The data of qRT PCR correlated with the RNA-seq results. All infected and treated samples showed elevated expression of the putative gene and (EUC67555.1) plant senescence-associated protein the treated samples showed high regulation of these genes (Fig 16). The Ribosomal protein L16 (chloroplast) (YP 009309192.1) and hypothetical protein AGABI2DRAFT 187811 (XP 006455547.1) genes showed similar results in terms of upregulation of gene expression. Other randomly chosen DEGs whose expression was investigated for downregulation include the sucrose transport protein SUC2, the zinc finger protein ZAT10-like, the inositol-3-phosphate synthase, and the protein NRT1/PTR FAMILY (KHN32802.1, XP 028211725.1, and XP 003551991.1). 2.13 (Fig 16). It was observed that the expression of DEGs examined by qRT PCR and RNA-seq data were both identical.

**Figure 16.**
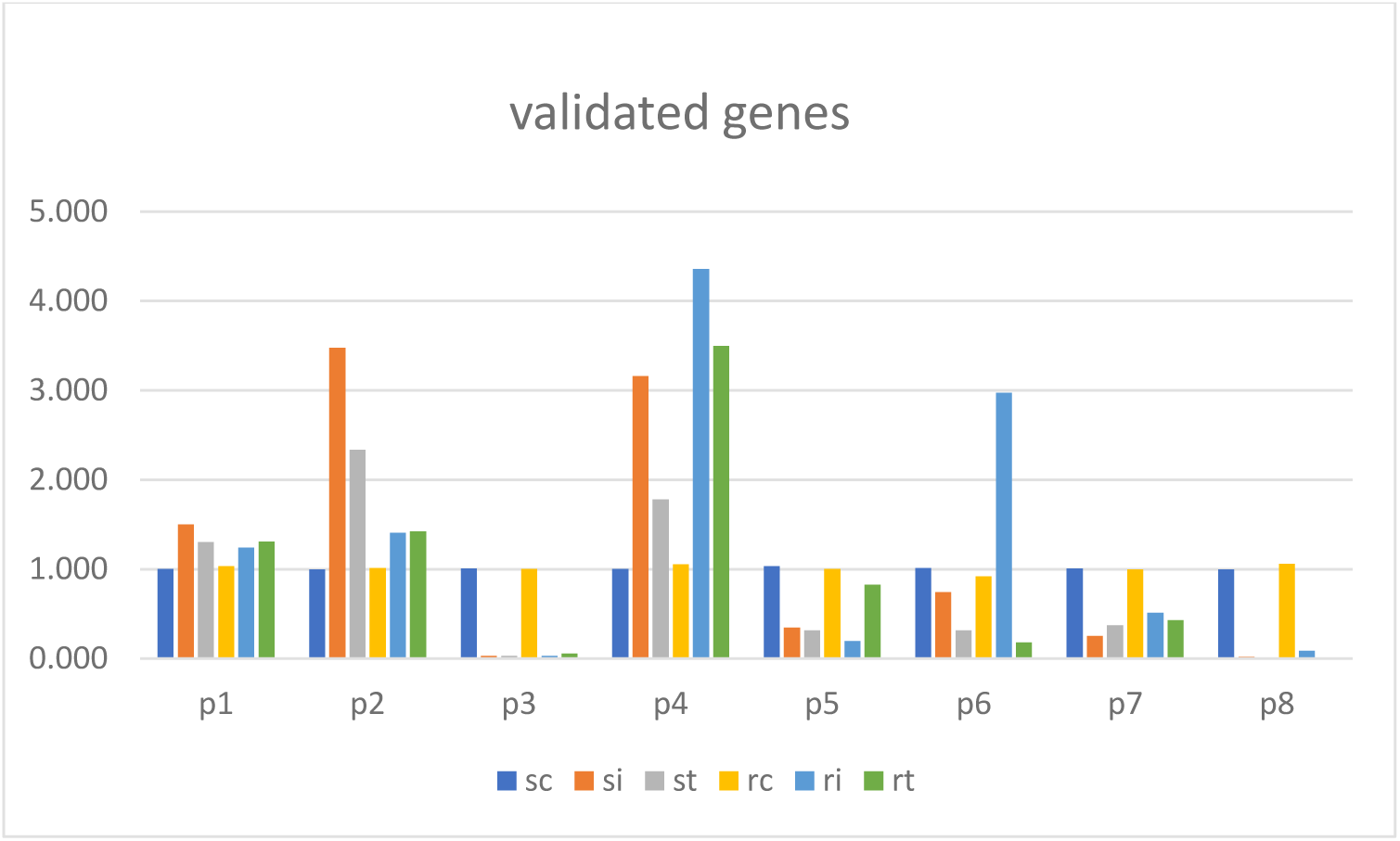
validation of differential expressed genes; sc: susceptible control, si: susceptible infected, st: susceptible treated, rc: resistant control; ri: resistant infected, rt: resistant treated, HKG: house kipping gene.

**Table 7:**
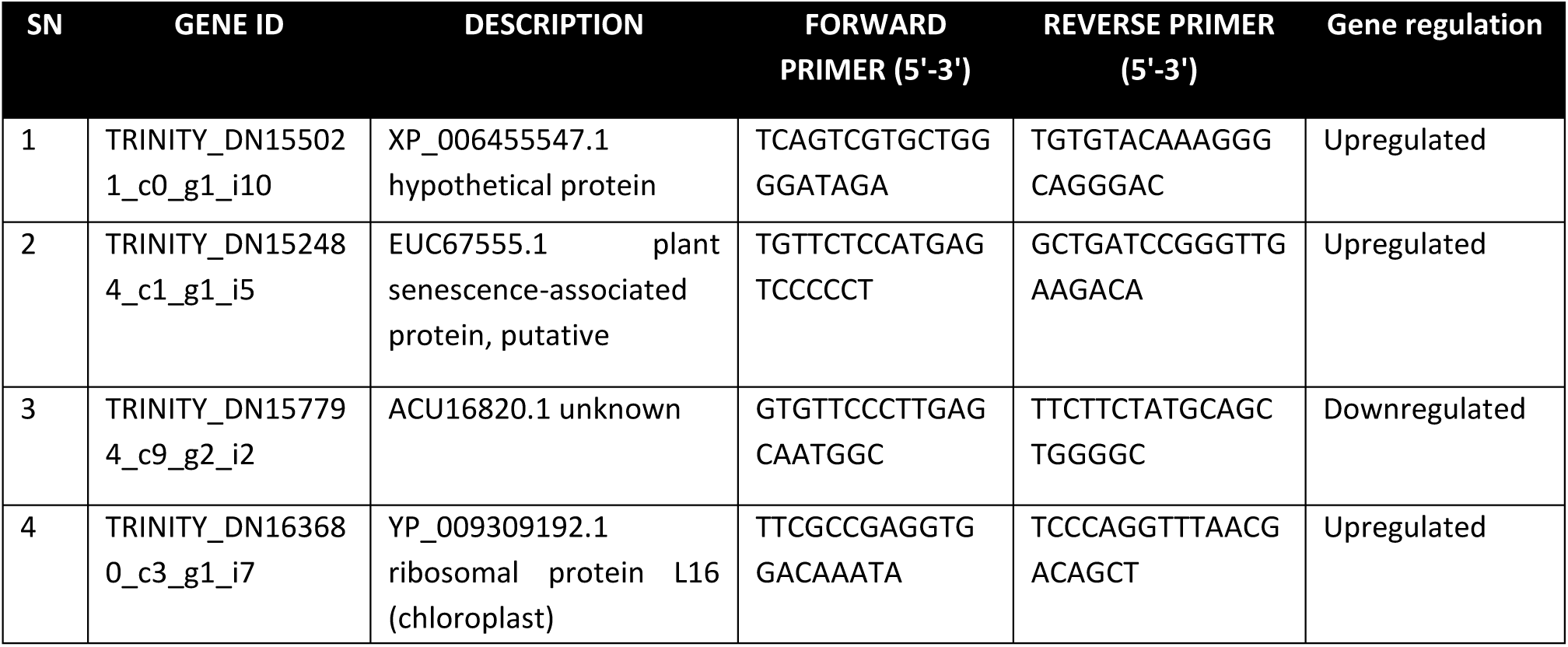
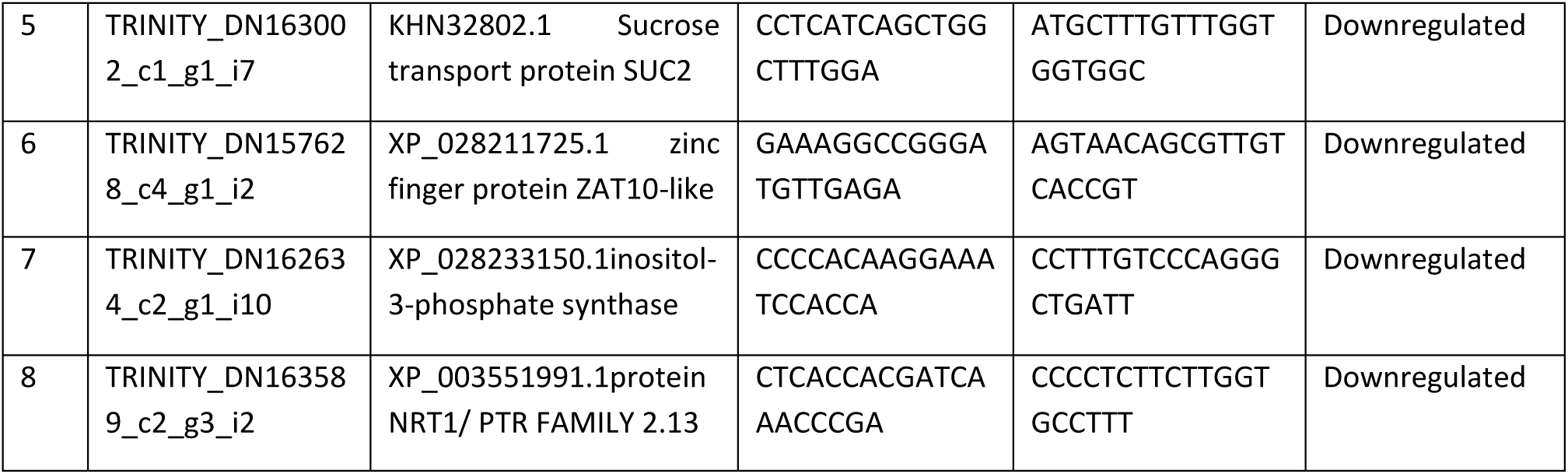
The molecular markers were designed for DEGs validation.

## 3. Discussion

In this study, a comprehensive transcriptomics analysis, coupled with supporting small-scale experiments, indicates that silicon treatment may confer protection to soybean plants against soybean *Macrophomina phaseolina* infection. By preventing the interaction between plant defense receptors and pathogen effectors. This is the first report on transcriptome analysis of the pathogenesis response to charcoal rot in two soybean genotypes, with or without silicon treatment. Many studies have shown that silicon can combat various pathogens [14, 15] but the exact role behind its infection prevention remain largely unanswered. Therefore, our research aims to fill this knowledge gap and provide the first RNA-Seq analysis of soybeans’ response to charcoal rot disease, particularly in the context of silicon treatment. Initial concept suggested that silicon’s protective role involved a mechanical barrier against fungal penetration, this hypothesis has waned in popularity due to evidence indicating that silicon application doesn’t notably increase leaf toughness. However, our study highlights a unique observation: soybean trichomes exhibited significant silicon accumulation, potentially contributing to enhanced leaf toughness and protection against fungal penetration. Previous study [16] made an intriguing discovery, they found that certain Arabidopsis plants, which couldn’t protect themselves from powdery mildew using a specific defense pathway, were still able to stay safe from this disease when they received silicon (Si). This suggests that silicon has a broader role than simply helping plants defend against diseases. When in present study soybean plants were treated with 1.7 mM Si, they showed strong protection against *Macrophomina phaseolina*. This was confirmed by the plant’s appearance and a heat map obtain after transcriptome analysis, which showed that the gene response in Si-treated infected plants was somewhat similar to that of healthy control plants. These results support the findings of scientist [17] demonstrated that plants treated with silicon (Si) didn’t exhibit a significant response to the presence of *P. sojae*.

Our data have conclusively demonstrated that treating soybean plants with silicon (Si) provides effective protection against *Macrophomina phaseolina*, which aligns with an earlier study [17]. The current study employed RNA-seq combined with potassium silicate supplementation to identify differentially expressed genes (DEGs) between various genotypes. The study uncovered a total of 3,106 DEGs, out of which 1,317 were found to be unique. Interestingly, the susceptible infection treatment exhibited the largest number of DEGs compared to the infection treatment in the resistant variety. This could be attributed to the resistant variety already expressing genes that confer resistance against *Macrophomina phaseolina*. The second highest number of DEGs was observed in the susceptible treated vs. resistant treated group, highlighting the effectiveness of the treatments and demonstrating the efficacy of silicon treatment. According to scientists [17] silicon (Si) has limited effects on healthy soybean plants. During *P. sojae* infection, Si influenced only around 50 out of 56,000 soybean genes, mostly downregulating them without specific pathway changes. This suggests that Si primarily helps plants by offering protection during times of stress. The current study’s findings are consistent with the findings of prior research.

A significant proportion of the differentially expressed genes (DEGs) identified in this study were associated with the ethylene-responsive factor (ERF), MYB, NAC, and FAR1 family proteins, indicating the presence of treatment-sensitive genes. The activation of ethylene pathways is well-documented in triggering immune responses to pathogen infections. Transcription factors (TFs) play a crucial role in the immune system by activating pathogenesis-related (PR) proteins and are closely linked to defense-related TF family members such as MYB, NAC, and ERF. Consistent with previous studies [18, 19], our research revealed elevated expression of ERF family proteins in resistant soybean genotypes, particularly the NP001235512 disease resistance protein. Other investigations have also demonstrated the induction of defense-related genes in response to pathogen infections. For instance, scientist [20] observed significant upregulation of callose synthase, PR genes, and receptor-like protein genes in *lomandra longifolia* roots infected with *Phytophinna momicnamomi*. Similarly, in apple root defense mechanisms against *F. solani* infection, reported differential expression of defense-related genes, including Cytochrome P450s and UDP-glycosyltransferases [12].

The present study observed an increase in the NAC family protein chitinase class-I precursor in both susceptible treated and resistant genotypes, indicating its involvement in resistance. Additionally, previous research has associated ERF/AP2, CDPK, and leucine-rich repeats with soybean resistance to charcoal rot [21, 22]. The NAC transcription factor family, known for its hypersensitivity to pathogen invasion, may activate chitinase to confer resistance. Moreover, the WRKY, NBs-LRR, and flavonoid pathways have been implicated in charcoal rot resistance [11, 23]. The scientist [11] investigated the ability of soybean to resist charcoal rot and reported the involvement of genes related to WRKY, NBs-LRR, and flavonoid pathways, which aligns with our findings. The WRKY protein, encoded by 174 genes and responsive to salicylic acid, contains zinc finger motifs and a conserved sequence [24], similar to the findings in our study where zinc finger proteins were significantly enhanced in both susceptible and resistant genotypes treated with silicon, along with the ribosomal-like protein L16. These observations suggest that zinc finger protein and ribosomal protein L16 may play a role in resistance mechanisms.

The current research KEGG enrichment analysis revealed a significant enrichment of ansamycin biosynthesis, which is known to promote plant growth under challenging conditions. Additionally, enhanced mechanisms for producing flavone and flavanol compounds were observed. The silicon treatment employed in present study induced defense responses, including the deposition of polyphenolic compounds, such as flavonoids, which play a crucial role in preventing pathogen growth. Flavonoids, similar to reactive oxygen species (ROS), function as stress protectors, while transcription factors (TFs) such as MYB and bHLH activate defense-related pathways [25–27]. The present research findings provide further support for the involvement of MYB, bHLH, and WD40 TFs in controlling the flavone and flavanol biosynthesis pathway, which exhibited higher expression levels in the resistant genotypes.

In the study at hand, histopathological analysis of the susceptible genotype’s roots infected with *Macrophomina phaseolina* revealed notable alterations in cell structure. The presence of the fungus’s mycelium caused obstruction within the vascular tissue of the affected roots, while the proliferation of microsclerotia became increasingly apparent over time. Importantly, we captured the dynamic process of hyphal penetration by *Macrophomina phaseolina* in real-time, providing valuable visual evidence of the infection process (https://youtube.com/shorts/t1iT5Hic2_M?feature=share). Additionally, the present study identified a genotype that exhibited significant suppression of hyphae penetration and multiplication, indicating its resistance to the pathogen. These findings align with the observations made by scientist [28] who reported similar outcomes in terms of epidermal penetration, hyphal development, and microsclerotia attachment to the root four days after inoculation.

The research sheds light on the molecular mechanisms underlying the interaction between *Macrophomina phaseolina* and soybean, which will aid in the development of strategies for breeding resistant plants. By utilizing the Illumina HiSeq platform and examining resistant and susceptible genes, we gained insight into the infection process of *Macrophomina phaseolina* in soybean genotypes. In the resistant genotype, we identified 41 defense-related genes, compared to only nine in the susceptible genotype. These findings provide a valuable resource for breeding *Macrophomina phaseolina* resistance, although further research is needed to elucidate the function of the identified resistant and susceptible genes.

## 4. Conclusion

In this study we find candidate genes related to susceptibility and resistance to *Macrophomina phaseolina* in soybean susceptible and resistant material. These findings provide give further insights regarding molecular mechanism underlying the interaction between host and pathogen and also effect of Si treatment in that scenario. Our results will be beneficial or helpful for resistant breeding practices. However, further reliable evidence is required to validate these results. For example, through VIGS transiently silencing genes to reveal their function will be helpful to validate these results and also use of genome editing via Cas9 to fully knockout the candidate gene will helpful to validate these results. This investigation, provide insight on the *Macrophomina phaseolina* infection process in soybean and revealed resistant and susceptible genes using Illumina sequencing platform and also the effect of silicate.

## 5. Materials and methods

### 5.1. Plant material and characterization of fungus

The experimental genotypes Suvarn soya (highly resistant) and TAMS-38 (highly susceptible) are developed by the Dr. Panjabrao Deshmukh Krishi Vidyapeeth, Akola through Mutation Breeding in collaboration with the Bhabha Atomic Research Center, Trombay. The pathogen *Macrophomina phaseolina* was taken from an infected soybean field in order to conduct a disease screening under *in vitro* condition. The pathogen was isolated using the serial dilution PDA method and microscopically validated using lactophenol cotton blue mounting solution. The genomic DNA was isolated from *Macrophomina phaseolina* using the DNAzol method [29], followed by ITS primer amplification, PCR product sequencing, BLAST analysis, and homology search. The sequence data submitted to NCBI for further reference and analysis.

### 5.2. On field screening of genotypes against charcoal rot disease

The genotypes were evaluated under sick plot at the Regional Research Center for Soybean, under Dr. Panjabrao Deshmukh Krishi Vidyapeeth, Amravati, Maharashtra State (20° 55’ 53.82” N. 77° 45’ 32.57” E). The disease incidence percentage was determined using the formula outlined [30]. The disease index was studied using a scale of 0-5 score.

Disease incidence (%) = Number of infected plants / Total number of plants × 100

### 5.3. In vitro screening: How does silicon respond to *Macrophomina phaseolina*?

Both susceptible and resistant genotypes were raised in pots, deliberately inoculated with a virulent strain of *Macrophomina phaseolina* (strain Akola-01), and kept in a climate-controlled greenhouse at a temperature and humidity of 30 to 35^0^C. Genotypes were screened using three different treatments of silicon (Si) and pathogen under *in vitro*. The treatments included as Control Treatment-1: Test genotypes raised in autoclaved soil without application of Si and *Macrophomina phaseolina*; Treatment-2 *Macrophomina phaseolina* treated: Pots inoculated with *M. phaseolina* without Si application; Treatment-3 Combined treatment of Si and *Macrophomina phaseolina*: Plants raised in pots treatment 1 and 2 irrigated with normal water and plants raised in treatment 3 pots were given weekly irrigations of 250 ml of a 1.7 mM potassium silicate. The plants were maintained under greenhouse until disease symptoms appeared and 42 days after sowing, leaf tissue samples were taken. The tissue samples from both test genotypes raised under different treatments were immediately frozen in liquid nitrogen and then preserved in RNA later at 4°C overnight. Following that, RNA-seq analysis was carried out using these samples.

### 5.4. Uptake and accumulation of K_2_SiO_3_

Tissue samples of susceptible and resistant genotypes raised under three treatments were subjected to scanning electron microscopy (SEM) and energy dispersive X-ray spectroscopy (EDX) analysis 45 days after seeding. Root and leaf tissues were examined using a Jeol JSM-IT300 SEM equipped with an EDX analyzer (EDAX, Octane Plus, Ametek, United States) to determine Si distribution. Si values were shown as weight percentages of the total Si element after the data was processed using the TEAM Enhanced ver. 4.3 program (EDAX Ametek, United States).

### 5.5. RNA sequencing

Six samples, comprising susceptible and resistant genotypes from all three treatments were collected from in-vitro experiment in three biological replications. RNA was then isolated using a plant RNA isolation kit, and 250 ng of total RNA was utilized for mRNA amplification using the NEB Next Poly (A) mRNA magnetic separation module. Illumina HiSeq was used to produce the sequencing data. The NEBNext Ultra-TM II RNA Library Prep Kit was used to prepare libraries from enriched mRNAs. Three biological replicates of each sample were used in the Illumina sequencing process.

#### 5.5.1. Transcriptome data processing

Raw reads were quality-checked and then subjected to data cleaning and pre-processing. The sequence data filtered using *Trimmomatic* v0.39 tool [31] by: 1) Removing reads containing a sequencing adapter; 2) Removing reads with a low-quality. Thus, clean reads/high quality reads were obtained and stored in FASTQC-0.11.8 tool [32]. De novo assembly of the transcriptome was performed with *Trinity* assembler v2.5.1 [33] and ‘Tuxedo2’ protocol (HISAT2 followed by StringTie2) was used for the reference-based assembly. The de novo assembly and reference assembly were compared for novel transcript using BLAST+ [34]. Reference assembly of *Glycine max* (Soybean) available at the ensemble was used as a reference model for the reference-based protocol (http://ftp.ensemblgenomes.org/pub/plants/release-53/fasta/glycine_max/dna/). The annotation file used for *Glycine max* is available at http://ftp.ensemblgenomes.org/pub/plants/release-53/gtf/glycine_max/. Keeping this in mind, the *do novo* assembly was used for the identification of novel differentially expressed transcripts, transcription factors, variants, and gene regulatory networks.

#### 5.5.2. Identification of differentially expressed genes (DEGs)

DEGs were identified using script available in Trinity tool. Transcript quantification was done using RSEM (RNA-Seq by expectation maximization) [35] to calculate the expression. This maps the clean reads onto the *de novo* transcriptomic assembly using Bowtie2 [36] perform the quantification using RSEM. In next step, DEGs were detected using the p value (<0.005) threshold in multiple tests and analyses was determined using FDR. The DEGs were deemed significant according to FDR<0.001, p-value <0.05 and log_2_fold change= ±2) the edgeR algorithm [37]. DEGs were identified in nine pairs of conditions i.e. (1) susceptible-control vs resistant-control (2) susceptible-control vs susceptible-infected (3) susceptible-control vs susceptible-treated (4) susceptible-infected vs resistant-infected (5) susceptible-treated vs resistant-treated (6) resistant-control vs resistant-infected (7) resistant-control vs resistant-treated (8) resistant-infected vs resistant-treated and (9) susceptible-infected vs susceptible-treated.

#### 5.5.3. Functional annotation, GO enrichment, and transcription factor identification

Functional annotation and GO annotation of obtained transcript was performed using Blast2GO Pro ver 3.1. Gene ontology (GO) enrichment analysis helps to understand the three important categories-biological processes, molecular functions and cellular components. KEGG (Kyoto Encyclopaedia of Genes and Genomes) pathways and enzyme classes were also identified [38]. KEGG pathway enrichment and GO enrichment were also performed to find the highly enriched KEGG pathways and GO terms [13]. For GO enrichment all the upregulated and downregulated DEGs from all the nine compared conditions were merged and then GO enrichment was performed using the Fisher’s Exact Test option available in Blast2Go software. KEGG pathways identified in DEGs were used for pathway enrichment analysis. BlastX search with E-value < 1e-03 was used for the identification of transcription factors (TFs) against the Plant TFDB for all the differential expressed genes obtained from nine combinations of datasets [39].

#### 5.5.4. Variant identification

Biological variations in biomarkers like SSR and nucleotide polymorphism were identified. Two types of biomarkers were identified *viz.* Simple Sequence Repeats (SSRs) and nucleotide polymorphism. SSRs were identified from the *de novo* assembled transcripts using MISA (MIcroSAtellite identification tool) to get the mono-, di-, tri, tetra-, Penta- and hexanucleotide and compound repeats [40]. Primers were also generated for SSRs using Primer3 [41] with parameters (annealing temperature-min: 57 °C, optimal: 60°C, maximum: 63 °C, primer size-min: 15, optimal: 18, maximum: 28 oligo-nucleotides).

Nucleotide polymorphisms like Single Nucleotide Polymorphism (SNPs), Multiple Nucleotide Polymorphism (MNPs), Insertion and Deletion (InDels), and complex structures were called using the Freebayes software [42]. Initially, the clean reads were mapped onto the transcriptome assembly using the Bowtie2 tool [36]. SAM tools was used to pre-process the alignment/map files (SAM/BAM) for sorting, duplicate removal, read group addition, and build the BAM index for the BAM file (Li, H. 2011). These indexed BAM files were further used in freebayes for variant calling. In order to obtain the significant variants, stringent parameters like minimum mapping quality ≥ 20 and minimum base quality > 20 were used.

#### 5.5.5. Gene regulatory networks

*Cytoscape* version 3.9.1 was used for the construction and visualization of gene regulatory networks (GRNs). ‘House Keeping Genes’, ‘Uncharacteristic proteins’ and ‘Similar genes were not considered in making the GRN. DEGs having high log fold change values were used for network construction by calculating the gene correlation using the Pearson Correlation Coefficient method. Normalized expression values were used for all nine conditions [44]. Hub genes were also identified after the identification of network centrality and topology and based on node degree.

#### 5.5.6. Soybean online database resource

For the availability of our research data to the scientific community, we have developed an online database resource for soybean. This database, (Soybean Transcriptome Database for Charcoal rot (STDbCr)) provides research data like transcripts annotations, DEGs, transcription factors, and biomarkers along with their primers and other genomic information in a single place. The database has been developed using the LAMP protocol (Linux Apache MySQL PHP). This resource is freely available for academic use at http://backlin.cabgrid.res.in/stdbcr.

#### 5.5.7. q-RT-PCR validation

Total RNA isolated and cDNA was synthesized using cDNA synthesis kit (Takara, Japan). The gene cons 15 (CDPK-related protein kinase) served as an internal control [45]. Primer sets were designed using primer3 software. The efficiency of the PCR was examined for primers that continuously indicated. The expression analysis was done by qRT-PCR (QIAquant Real-Time Thermal Cycler) conducted at ICAR-DFR, Pune institute. q-RT-PCR was done with FastStart SYBR green master mix. Using ΔΔCT, the expression analysis was calculated and represented as fold increase= 2^−ΔΔCT^ [46].

## Supplementary Information (Starting from this point, the article comprises more than 2000 words.)

- Additional file_Figure_1: Average trend of 3 days interval progress of Charcoal rot disease in soybean genotypes
- Additional file_video_1: The live movement of *Macrophomina phaseolina* in soybean root, time-laps video for 4-5hrs.
- Additional file_Figure_3: Comparative study of Scanning Electron Micrographs and EDX analysis for uptake and accumulation of silicon in soybean genotype cv., TAMS-38, tissue leaf and root.
- Supplementary Figure 1: Heatmap illustrating Differentially Expressed Genes (DEGs)
- Supplementary Figure 2: MA-Volcano plot representing Differentially Expressed Genes (DEGs)
- Supplementary Figure 3: Distribution of the top-hit species identified in the dataset, showing the relative abundance of sequence hits assigned to different taxonomic categories.
- Supplementary_Sheet_1_DEGs_in_9_comparisons
- Supplementary_Sheet_2_GO_of_idnetified_DEGs
- Supplementary_Sheet_3_GO_enrichment
- Supplementary_Sheet_4_KEGG_Pathway in DEGs
- Supplementary_Sheet_5_Identified_TF_in_DEGs
- Supplementary_Sheet_6_Identified SSR and primers
- Supplementary_Sheet_7_Identified variants in assembly
- Supplementary_Sheet_8_DEGs_used_for_GRN

## Acknowledgements

The authors express their gratitude for the support received from Dr. Panjabrao Deshmukh Krishi Vidyapeeth, Akola, Maharashtra State, and the Department of Biotechnology (DBT), India, which supported to this study. Additionally, the authors extend their appreciation to the laboratory members for their valuable technical assistance and insightful suggestions during the manuscript preparation.

## Author contributions

P. V. J. conceptualized and designed the study; S. G. M. and P. K. S. wrote the original draft of the manuscript. A field experiment was conducted and carried out under sick plot and pot conditions by S. G. M., S. S. N., R. S. G., E. R. V., and M. P. M. A gene validation experiment was carried out by S. G. M., P. G. K., and P. R. J., S. J. and M. A. I. completed the computational analysis. S. S. M., S. B. S., R. B. G., R. D., H. S. annotated genes as well as performed functional gene classification and differentials gene expression analysis and reviewed the draft manuscript; D. K. and V. K. K. reviewed the analysis and critically reviewed the final draft manuscript. P. V. J. was responsible for funding acquisition. All authors read and approved the final manuscript.

## Funding

The study was supported by a Dr. Panjabrao Deshmukh Krishi Vidyapeeth, Akola, Maharashtra State and Department of Biotechnology (DBT), India.

## Availability of data and material

▪ A website resource (STDbCr) has been created for soybean for the general public to utilize (http://backlin.cabgrid.res.in/stdbcr/).
▪ The RNAseq data for the study has been deposited in a repository with the BioProject title “The RNAseq Study on Charcoal Rot Fungal Disease in Soybean.” This project has been assigned the accession number PRJNA1026744. The data for this study can be accessed through the following link: (https://www.ncbi.nlm.nih.gov/bioproject/?term=PRJNA1026744)
▪ The sequence for *Macrophomina phaseolina* has been deposited in the repository with the GenBank accession number MZ823608.1. The sequence can be access using the following link: (https://www.ncbi.nlm.nih.gov/search/all/?term=MZ823608.1.)
▪ A live movement of *Macrophomina phaseolina* in soybean root; link for same video. (https://youtube.com/shorts/t1iT5Hic2_M?feature=share).

## Declarations

## Conflicts of interest

The authors declare that they have no competing interest.

## Ethics approval and consent to participate

Not applicable.

## Adherence to national and international regulations

Not applicable.

## Consent for publication

Not applicable.

## Competing interests

The authors declare that they have no competing interests.

## Notes

### Competing Interest Statement

The authors have declared no competing interest.

## Reference

1. Jaiswal S, Jadhav P V., Jasrotia RS, Kale PB, Kad SK, Moharil MP, et al. Transcriptomic signature reveals mechanism of flower bud distortion in witches’-broom disease of soybean (Glycine max). BMC Plant Biol. 2019;19.

2. Shoaib A, Ali H, Javaid A, Awan ZA. Contending charcoal rot disease of mungbean by employing biocontrol Ochrobactrum ciceri and zinc. Physiology and Molecular Biology of Plants. 2020;26.

3. Sarr MP, Ndiaye M, Groenewald JZ, Crous PW. Genetic diversity in *Macrophomina phaseolina*, the causal agent of charcoal rot. Phytopathol Mediterr. 2014;53.

4. Kaur S, Dhillon GS, Brar SK, Vallad GE, Chand R, Chauhan VB. Emerging phytopathogen *Macrophomina phaseolina*: Biology, economic importance and current diagnostic trends. Critical Reviews in Microbiology. 2012;38.

5. Mengistu A, Ray JD, Smith JR, Arelli PR, Bellaloui N, Chen P, et al. Effect of charcoal rot on selected putative drought tolerant soybean genotypes and yield. Crop Protection. 2018;105.

6. da Silva MP, Klepadlo M, Gbur EE, Pereira A, Mason RE, Rupe JC, et al. QTL mapping of charcoal rot resistance in PI 567562A soybean accession. Crop Sci. 2019;59:474–9.

7. Coskun D, Deshmukh R, Sonah H, Menzies JG, Reynolds O, Ma JF, et al. The controversies of silicon’s role in plant biology. New Phytologist. 2019;221.

8. Hussain S, Mumtaz M, Manzoor S, Shuxian L, Ahmed I, Skalicky M, et al. Foliar application of silicon improves growth of soybean by enhancing carbon metabolism under shading conditions. Plant Physiology and Biochemistry. 2021;159.

9. Mandlik R, Thakral V, Raturi G, Shinde S, Nikolić M, Tripathi DK, et al. Significance of silicon uptake, transport, and deposition in plants. Journal of Experimental Botany. 2020;71.

10. Sahebi M, Hanafi MM, Siti Nor Akmar A, Rafii MY, Azizi P, Tengoua FF, et al. Importance of silicon and mechanisms of biosilica formation in plants. BioMed Research International. 2015;2015.

11. Deshmukh R, Tiwari S. Molecular interaction of charcoal rot pathogenesis in soybean: a complex interaction. Plant Cell Reports. 2021;40.

12. Xiang L, Wang M, Pan F, Wang G, Jiang W, Wang Y, et al. Transcriptome analysis Malus domestica ‘M9T337’ root molecular responses to Fusarium solani infection. Physiol Mol Plant Pathol. 2021;113.

13. Kanehisa M, Goto S. KEGG: Kyoto Encyclopedia of Genes and Genomes. Nucleic Acids Res. 28, 27-30 (2000). Nucleic Acids Res. 2000;28:27–30.

14. Fauteux F, Rémus-Borel W, Menzies JG, Bélanger RR. Silicon and plant disease resistance against pathogenic fungi. FEMS Microbiology Letters. 2005;249:1–6.

15. Bélanger RR, Benhamou N, Menzies JG. Biochemistry and Cell Biology Cytological Evidence of an Active Role of Silicon in Wheat Resistance to Powdery Mildew (Blumeria graminis f. sp. tritici). 2003.

16. Vivancos J, Labbé C, Menzies JG, Bélanger RR. Silicon-mediated resistance of Arabidopsis against powdery mildew involves mechanisms other than the salicylic acid (SA)-dependent defence pathway. Mol Plant Pathol. 2015;16:572–82.

17. Rasoolizadeh A, Labbé C, Sonah H, Deshmukh RK, Belzile F, Menzies JG, et al. Silicon protects soybean plants against Phytophthora sojae by interfering with effector-receptor expression. BMC Plant Biol. 2018;18.

18. Amorim L, Santos R, Neto J, Guida-Santos M, Crovella S, Benko-Iseppon A. Transcription Factors Involved in Plant Resistance to Pathogens. Curr Protein Pept Sci. 2016;18.

19. Rong W, Qi L, Wang A, Ye X, Du L, Liang H, et al. The ERF transcription factor TaERF3 promotes tolerance to salt and drought stresses in wheat. Plant Biotechnol J. 2014;12:468–79.

20. Dong X, Hong Z, Chatterjee J, Kim S, Verma DPS. Expression of callose synthase genes and its connection with Npr1 signaling pathway during pathogen infection. Planta. 2008;229:87–98.

21. Coser SM, Reddy RVC, Zhang J, Mueller DS, Mengistu A, Wise KA, et al. Genetic architecture of charcoal rot (*Macrophomina phaseolina*) resistance in soybean revealed using a diverse panel. Front Plant Sci. 2017;8.

22. Yuan X, Wang H, Cai J, Li D, Song F. NAC transcription factors in plant immunity. Phytopathology Research. 2019;1.

23. Radadiya N, Mangukia N, Antala V, Desai H, Chaudhari H, Dholaria TL, et al. Transcriptome analysis of sesame-*Macrophomina phaseolina* interactions revealing the distinct genetic components for early defense responses. Physiology and Molecular Biology of Plants. 2021;27.

24. Yang Y, Zhou Y, Chi Y, Fan B, Chen Z. Characterization of Soybean WRKY Gene Family and Identification of Soybean WRKY Genes that Promote Resistance to Soybean Cyst Nematode. Sci Rep. 2017;7.

25. August PR, Tang L, Yoon YJ, Ning S, Müller R, Yu TW, et al. Biosynthesis of the ansamycin antibiotic rifamycin: Deductions from the molecular analysis of the rif biosynthetic gene cluster of Amycolatopsis mediterranei S699. Chem Biol. 1998;5.

26. Ahammed GJ, Yang Y. Mechanisms of silicon-induced fungal disease resistance in plants. Plant Physiology and Biochemistry. 2021;165:200–6.

27. Ahammed GJ, Yang Y. Anthocyanin-mediated arsenic tolerance in plants. Environmental Pollution. 2022;292.

28. Hemmati P, Zafari D, Mahmoodi SB, Hashemi M, Gholamhoseini M, Dolatabadian A, et al. Histopathology of charcoal rot disease (*Macrophomina phaseolina*) in resistant and susceptible cultivars of soybean. Rhizosphere. 2018;7.

29. Sinha R, Irulappan V, Mohan-Raju B, Suganthi A, Senthil-Kumar M. Impact of drought stress on simultaneously occurring pathogen infection in field-grown chickpea. Sci Rep. 2019;9.

30. Irulappan V, Mali KV, Patil BS, Manjunatha H, Muhammad S, Senthil-Kumar M. A sick plot–based protocol for dry root rot disease assessment in field-grown chickpea plants. Appl Plant Sci. 2021;9.

31. Bolger AM, Lohse M, Usadel B. Trimmomatic: A flexible trimmer for Illumina sequence data. Bioinformatics. 2014;30.

32. Andrews S. FastQC - A quality control tool for high throughput sequence data. http://www.bioinformatics.babraham.ac.uk/projects/fastqc/. Babraham Bioinformatics. 2010.

33. Haas BJ, Papanicolaou A, Yassour M, Grabherr M, Blood PD, Bowden J, et al. De novo transcript sequence reconstruction from RNA-seq using the Trinity platform for reference generation and analysis. Nat Protoc. 2013;8.

34. Camacho C, Coulouris G, Avagyan V, Ma N, Papadopoulos J, Bealer K, et al. BLAST+: Architecture and applications. BMC Bioinformatics. 2009;10.

35. Li B, Dewey CN. RSEM: Accurate transcript quantification from RNA-Seq data with or without a reference genome. BMC Bioinformatics. 2011;12.

36. Langmead B, Salzberg SL. Fast gapped-read alignment with Bowtie 2. Nat Methods. 2012;9.

37. Robinson MD, McCarthy DJ, Smyth GK. edgeR: A Bioconductor package for differential expression analysis of digital gene expression data. Bioinformatics. 2009;26.

38. Conesa Stefan A and G. International Journal of Plant Genomics. Bioinformatics Tools for Plant Genomics. 2008;2008.

39. Jin X, Huang H, Wang L, Sun Y, Dai S. Transcriptomics and metabolite analysis reveals the molecular mechanism of anthocyanin biosynthesis branch pathway in different senecio cruentus cultivars. Front Plant Sci. 2016;7 September.

40. Thiel T, Michalek W, Varshney RK, Graner A. Exploiting EST databases for the development and characterization of gene-derived SSR-markers in barley (Hordeum vulgare L.). Theoretical and Applied Genetics. 2003;106.

41. Untergasser A, Cutcutache I, Koressaar T, Ye J, Faircloth BC, Remm M, et al. Primer3-new capabilities and interfaces. Nucleic Acids Res. 2012;40.

42. Garrison, E., & Marth G. Haplotype-based variant detection from short-read sequencing. Journal of Clinical Gastroenterology. 2012;51.

43. Li. A statistical framework for SNP calling, mutation discovery, association mapping and population genetical parameter estimation from sequencing data. Bioinformatics. 2011;27.

44. Shannon P, Markiel A, Ozier O, Baliga NS, Wang JT, Ramage D, et al. Cytoscape: A software Environment for integrated models of biomolecular interaction networks. Genome Res. 2003;13.

45. Libault M, Thibivilliers S, Bilgin DD, Radwan O, Benitez M, Clough SJ, et al. Identification of Four Soybean Reference Genes for Gene Expression Normalization. Plant Genome. 2008;1.

46. Livak KJ, Schmittgen TD. Analysis of relative gene expression data using real-time quantitative PCR and the 2-ΔΔCT method. Methods. 2001;25.

